# A Genome-wide Library of MADM Mice for Single-Cell Genetic Mosaic Analysis

**DOI:** 10.1101/2020.06.05.136192

**Authors:** Ximena Contreras, Amarbayasgalan Davaatseren, Nicole Amberg, Andi H. Hansen, Johanna Sonntag, Lill Andersen, Tina Bernthaler, Anna Heger, Randy Johnson, Lindsay A. Schwarz, Liqun Luo, Thomas Rülicke, Simon Hippenmeyer

## Abstract

Mosaic Analysis with Double Markers (MADM) offers a unique approach to visualize and concomitantly manipulate genetically-defined cells in mice with single-cell resolution. MADM applications include the analysis of lineage; single-cell morphology and physiology; genomic imprinting phenotypes; and dissection of cell-autonomous gene functions *in vivo* in health and disease. Yet, MADM could only be applied to <25% of all mouse genes on select chromosomes thus far. To overcome this limitation, we generated transgenic mice with knocked-in MADM cassettes near the centromeres of all 19 autosomes and validated their use across organs. With this resource, >96% of the entire mouse genome can now be subjected to single-cell genetic mosaic analysis. Beyond proof-of-principle, we applied our MADM library to systematically trace sister chromatid segregation in distinct mitotic cell lineages. We found striking chromosome-specific biases in segregation patterns, reflecting a putative mechanism for the asymmetric segregation of genetic determinants in somatic stem cell division.

## INTRODUCTION

Genetic mosaic individuals contain cells of distinct genotypes. The phenomenon of genetic mosaicism occurs naturally and is widespread across multicellular organisms. Mosaicism may progressively emerge during life but remain silent with no obvious or severe phenotypic consequences for extended periods of time (Yizhak et al. 2019). However, mosaicism is also associated with a number of pathologies in human including cancer which emerges as a clone of one mutated single cell, or many neurological disorders affecting brain development and function (Biesecker and Spinner 2013; D’Gama and Walsh 2018). Genetic mosaic animals have been experimentally created in a number of species including *C. elegans*, *Drosophila*, and mice among others; and such mosaic analyses provided many fundamental insights in a variety of biological systems (Germani et al. 2018; Kim et al. 2019; Lee and Luo 1999, 2001; Lozano and Behringer 2007; Luo 2007; Rossant and Spence 1998; Xu and Rubin 1993; Yochem and Herman 2003; Zong et al. 2005; Zugates and Lee 2004).

One of the most powerful applications inherent to induced genetic mosaics is the ability to alter gene function at high spatiotemporal resolution. In other words, a certain tissue can contain homozygous mutant cells for a gene of interest and wild-type cells in the same animal whose phenotypes can be compared with each other directly. If the genetic mosaic is sparse, even essential genes can be manipulated without affecting the overall health or viability of the animal. Furthermore, sparse genetic mosaics provide a highly effective means to study the causal relationship of genetic alteration and phenotypic manifestation at the individual cell level. Genetic mosaics also facilitate the analysis of cell competition and provide an assay to create models of disease. Historically, for over a century and up to date genetic mosaics have been most extensively generated and used in the fruit fly by capitalizing upon mitotic recombination between homologous chromosomes (Hotta and Benzer 1970; Lee and Luo 1999, 2001; Morgan and Bridges 1919; Stern 1936; Xu and Rubin 1993; Zugates and Lee 2004). Although technically slightly more challenging, the generation of genetic mosaics in mice - one of the most widely used model organisms to study gene function in health and disease - is becoming routine. A number of experimental approaches have been established including Mosaic Analysis with Double Markers (MADM) which is also based on mitotic recombination (Hippenmeyer 2013; Luo 2007; Tasic et al. 2012; Zong et al. 2005).

MADM relies on Cre/LoxP-mediated interchromosomal recombination to simultaneously generate homozygous mutant cells for a candidate gene of interest and wild-type cells in an otherwise heterozygous background. The induction of genetic mosaicism can be spatiotemporally controlled in genetically-defined cell-types by the use of tissue-specific Cre/ER-driver lines. Concurrent to the generation of genetic mosaicism, two split genes, encoding green fluorescent protein (GFP) and tdTomato (tdT) fluorescent markers, are reconstituted permitting unequivocal tracing of individual cellular phenotypes in mutant, heterozygous and wild-type cells, each labelled in distinct colors with 100% accuracy (Hippenmeyer et al. 2010; Zong et al. 2005) (Figure 1A; Figure S1).

**Figure 1.**
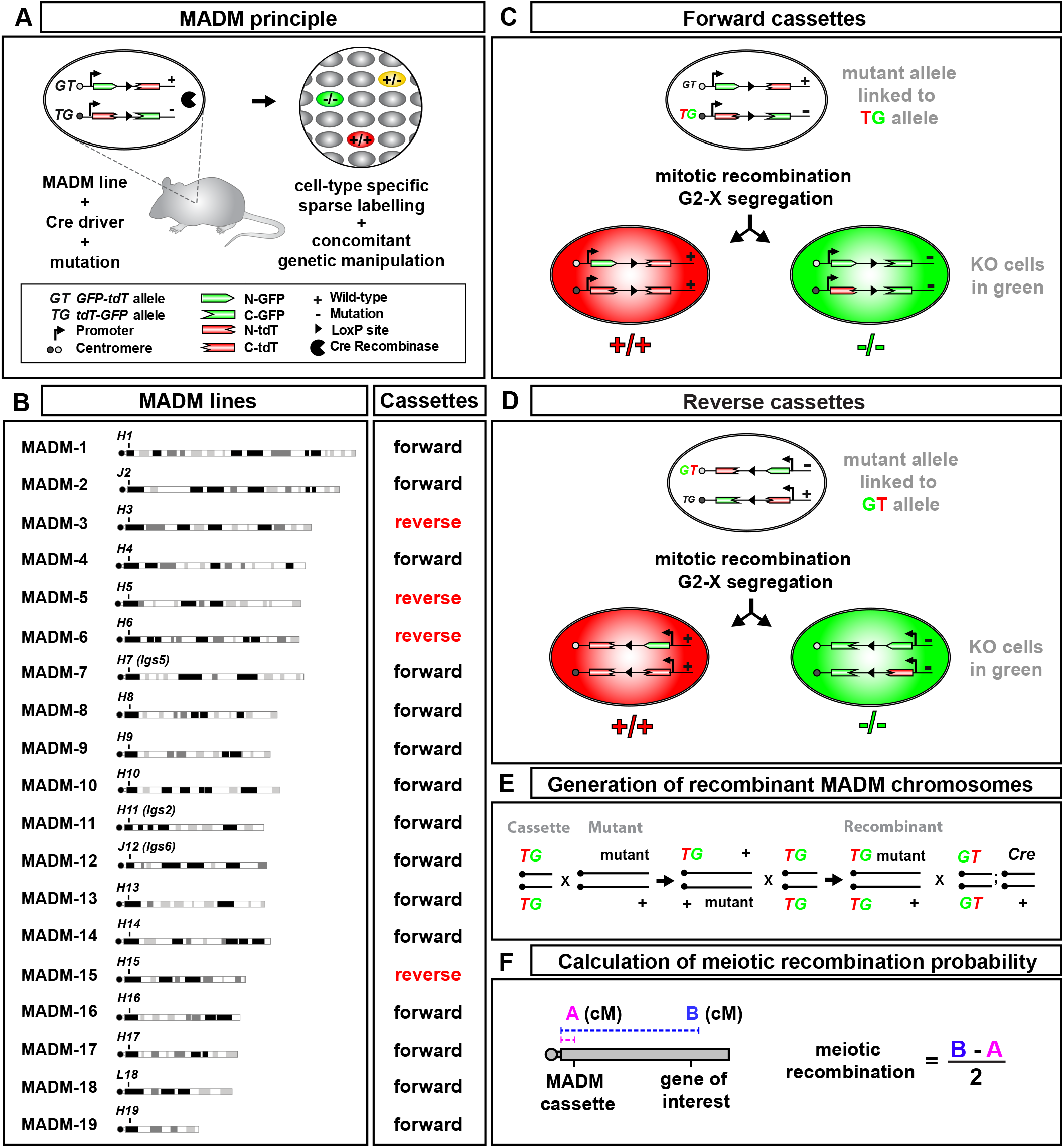
Extension of MADM to all 19 Mouse Autosomes. **(A)** Summary of the MADM principle. MADM enables the concomitant fluorescent labeling and genetic manipulation of genetically-defined cellular populations at clonal and single cell resolution. For MADM, two chimeric split marker genes containing partial coding sequences of eGFP and tdT are inserted into identical genomic loci of homologous chromosomes. Following Cre recombinase-mediated interchromosomal (trans) recombination during mitosis the split marker genes are reconstituted and functional green and red fluorescent proteins expressed. As a result, green GFP^+^, red tdT^+^ and yellow GFP^+^/tdT^+^ cells appear sparsely, due to inherently low stochastic interchromosomal recombination rate, within the genetically-defined cell population expressing Cre recombinase. Introduction of a mutant allele distal to the MADM cassette results in a genetic mosaic with homozygous mutant cells labeled in one color (eg. green GFP^+^) and homozygous wild-type sibling cells in the other (eg. red tdT^+^). Heterozygous cells appear in yellow (GFP^+^/tdT^+^). **(B)** Expansion of MADM to all mouse autosomes. Transgenic mice with MADM cassettes inserted close to the centromere have been for all 19 mouse autosomes. The directionality (forward, centromere-telomere and reverse, telomere-centromere) of marker gene transcription is indicated. **(C)** MADM labeling scheme for cassettes inserted in forward direction. MADM experiments involving forward cassettes require that the mutant allele of a candidate gene must be linked to the T-G MADM cassette in order for mutant cells to be labelled in green upon a G2-X MADM event. **(D)** MADM labeling scheme for cassettes inserted in reverse direction. MADM experiments involving reverse cassettes require that the mutant allele of a candidate gene must be linked to the G-T MADM cassette in order for mutant cells to be labelled in green upon a G2-X MADM event. **(E)** Generation of recombinant MADM chromosomes. To genetically link a mutant allele of a candidate gene of interest to the corresponding chromosome containing the T-G MADM cassette (i.e. forward orientation) it is necessary to first cross mice bearing the T-G MADM cassette with mice bearing the mutant allele. Resulting F1 transheterozygous offspring are then backcrossed to mice homozygous for T-G MADM cassette. In F2 recombinant offspring emerge from meiotic recombination events in the germline. These F2 recombinants now contain both the MADM cassette (in homozygous configuration) and the mutant allele linked on the same chromosome. For experimental MADM mice F2 recombinants are crossed with mice bearing G-T MADM cassette and a Cre driver of interest. **(F)** Calculation of predicted meiotic recombination frequency. The probability for meiotic recombination resulting in the linkage of the MADM cassette with the mutant allele can be estimated by the genetic distance of the MADM cassette to the location of the mutant allele divided by two.

One application of the MADM technology is lineage tracing - analysis of cell division patterns and distribution of clonally-related cells that constitute different parts of an organ. The temporally controlled induction of MADM with its dual marker property provides exact and unique information not only about birth dates of clones but also regarding their cell division patterns. Thus, MADM has been frequently utilized in the past to study the proliferation behavior of progenitor stem cells in a variety of tissues - including embryonic and adult neural stem cells (Beattie et al. 2017; Bonaguidi et al. 2011; Espinosa and Luo 2008; Gao et al. 2014; Kaplan et al. 2017; Liang et al. 2013; Llorca et al. 2019; Lv et al. 2019; Mayer et al. 2015; Mihalas and Hevner 2018; Ortiz-Alvarez et al. 2019; Picco et al. 2019; Shi et al. 2017; Wong et al. 2018); cardiomyocyte-(Ali et al. 2014; Devine et al. 2014; Mohamed et al. 2018) and pancreatic progenitor cells (Brennand et al. 2007; Desgraz and Herrera 2009; Salpeter et al. 2010); progenitors in the developing kidney (Riccio et al. 2016); and mesenchymal progenitors in the developing lung (Kumar et al. 2014). MADM enables high resolution single cell/clonal labeling and permits tracing of complex morphogenetic processes in 4D by using live-imaging over prolonged periods besides analysis of static time points (Hippenmeyer et al. 2010; Riccio et al. 2016).

MADM technology has recently emerged as a unique approach to probe genomic imprinting and the function of individual imprinted genes (Hippenmeyer et al. 2013; Laukoter et al. 2020). Genomic imprinting is a prevalent epigenetic phenomenon in placental mammals and results in the preferential expression of either the maternal or paternal inherited allele of a subset of genes (Barlow and Bartolomei 2014; Ferguson-Smith 2011; Tucci et al. 2019). MADM can be applied to create uniparental chromosome disomy (UPD, somatic cells with two copies of either the maternal or paternal chromosome) and visualize imprinting effects at morphological and transcriptional level with single cell resolution (Hippenmeyer et al. 2013; Laukoter et al. 2020).

Perhaps the most salient and unique property of the MADM technology is to create genetic mosaicism and thus conditional gene knockouts in single cells with 100% correlation between fluorescent labeling and genetic alteration (Beattie et al. 2017; Espinosa et al. 2009; Gao et al. 2014; Henderson et al. 2019; Hippenmeyer et al. 2010; Joo et al. 2014; Liang et al. 2013; Lv et al. 2019; Ortiz-Alvarez et al. 2019; Riccio et al. 2016). MADM-labeled wild-type and mutant cells in mosaic mice can be directly assessed by histological means, physiological analysis, and optical imaging *in vivo*. One clinically-relevant application of MADM is the tracing of tumor growth upon ablation of tumor suppressor genes in a small subset of cells in a particular tissue. Cells that are homozygous mutant for a candidate tumor suppressor gene are uniquely labeled and can be followed dynamically *in vivo* to study tumor progression and metastasis and/or to assay for the effects of therapeutic agents. As such, MADM has been used for the analysis of tumor formation and the delineation of cancer cell of origin at single cell level in the brain and distinct organs (Gonzalez et al. 2018; Liu et al. 2011; Muzumdar et al. 2016; Muzumdar et al. 2007; Yao et al. 2020). MADM could in principle be exploited to systematically study the loss of any tumor suppressor in the entire genome and for identifying the cellular origins of a wide variety of cancers.

The single cell phenotype in classic conditional or full knockout mutants often reflects a combination of both cell-autonomous gene function and environment-derived cues which may remedy or exacerbate any observed phenotype. It is thus important in genetic loss of function models to qualitatively and quantitatively determine the relative contributions of the intrinsic and extrinsic components to the overall loss of function phenotype. To this end, the MADM system offers an unmatched experimental solution. The candidate gene function can be either ablated in just very sparse mosaic and/or single clones (see above), or tissue-wide in all cells. However in both paradigms, single cell MADM labeling enables the high resolution quantitative phenotypic analysis (Beattie et al. 2017; Joo et al. 2014; Laukoter et al. 2020). The above paradigms thus potentially permit systematic dissection of the level of cell-autonomy of any gene function in a given tissue, provided appropriate Cre-driver lines exist.

A major current limitation of the MADM technology is that it can only be applied to study candidate genes located on Chr. 7, Chr. 11, Chr. 12, and distal to the *Rosa26* locus on Chr. 6, where MADM cassettes have been introduced (Zong et al., 2005; Hippemmeyer et al. 2010; Hippmeneyer et al., 2013). Thus, less than 25% of all genes in the mouse genome can be subjected to MADM analysis as described above. Here we overcome this constraint and expanded MADM technology to all mouse autosomes. We provide validation of all MADM reporters and quantitative assessment of the efficacy of MADM labeling in a variety of organs, tissues and a number of clinically-relevant stem cell niches across the entire mouse. Furthermore, we utilized the newly engineered MADM chromosomes to systematically determine sister chromatid segregation patterns in several somatic cell lineages. Our analysis revealed for the first time *in vivo* that sister chromatid segregation patterns in mitotic progenitor cell divisions are highly biased in a chromosome-specific manner, and are further affected by cell type.

## RESULTS

### Expansion of MADM to all Mouse Autosomes

For MADM, two reciprocally chimeric marker genes need to be targeted to identical loci on homologous chromosomes (Zong et al. 2005). The chimeric marker genes (*GT* and *TG* alleles) consist of N- or C-terminal halves of the coding sequences for green (enhanced green fluorescent protein, [G]) and red (tdTomato, [T]) fluorescent proteins interspersed by an intron with the loxP site (Hippenmeyer et al. 2010) (Figure 1A, Figure S1). Here we expand MADM to all nineteen mouse autosomes with the goal to enable MADM for the vast majority, nearly genome-wide, of autosomal genes in the mouse genome. Mouse autosomes consist of only one chromosome arm (i.e. telocentric conformation). We thus rationalized that inserting the MADM cassettes as close as possible to the centromere would maximize the number of genes located distally to the MADM cassette insertion site for prospective MADM experiments (Hippenmeyer et al. 2013; Hippenmeyer et al. 2010) (Figure 1A and 1B).

To identify suitable genomic loci for MADM cassette targeting, we applied a number of key criteria besides the one of closest possible distance to the centromere (Hippenmeyer et al. 2010). As such, the targeting loci should 1) locate to intergenic regions to minimize the probability of disrupting endogenous gene function; and 2) permit spatially and temporally ubiquitous and biallelic expression of the reconstituted GFP and tdT markers. To fulfill the first criteria we mapped gene-by-gene the genetic landscape of the centromeric-most 20 Mbp of all autosomes using the UCSC Genome Browser (www.genome.ucsc.edu; GRCm38/mm10); except Chr 7, 11 and 12 which previously have been rendered MADM-ready by employing the above criteria (Hippenmeyer et al. 2013; Hippenmeyer et al. 2010). Next we anticipated that the broadness of spatial and temporal EST (expression sequence tag) expression patterns of the neighboring genes flanking the putative targeting site would serve as proxy for the spatiotemporal extent of transgene expression. The final choice of the prime targeting loci (Figure 1B, Figure S2, Table 1) was based upon the most ideal combination of the three above key criteria. In total, more than 20’000 protein-coding genes, corresponding to >96% of the entire annotated mouse genome (GRCm38/mm10), are located distally to the MADM targeting loci across all 19 autosomes (Table 1).

**Table 1.**
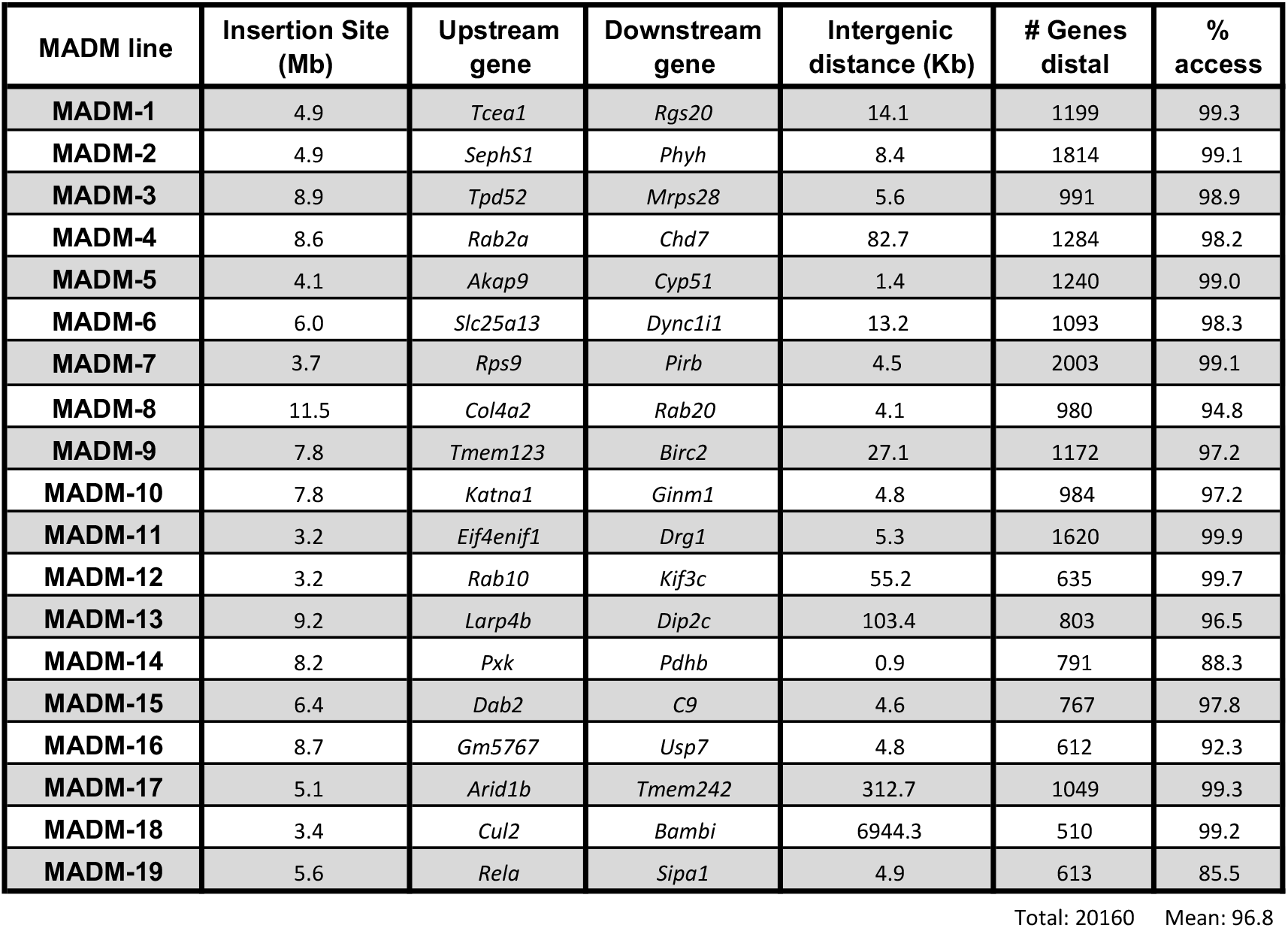
Location of Genomic Targeting Sites for MADM Cassette Insertion. Table lists the insertion sites, genes flanking the MADM cassette insertion site, distance between neighboring genes, number of genes located distally to the MADM cassettes, and percentage of access of genes that can be subjected to MADM analysis in all 19 MADM lines. In total 20’160 protein coding mouse genes, or 96.8% of them, are located distally to the MADM cassettes.

Next we cloned the selected genomic targeting loci, and inserted the MADM cassettes (Hippenmeyer et al. 2010) by homologous recombination in mouse ES cells (Figure S3, see STAR Methods for details). For most chromosomes, MADM cassettes were inserted in centromere-to-telomere transcriptional direction (Figure 1B, forward) except for Chr. 3, Chr. 5, Chr. 6, and Chr. 15 which required opposite directionality (Figure 1B, reverse) in order to best fulfill our locus choice criteria. The direction of reconstituted MADM marker gene transcription, upon interchromosomal recombination, has consequences for the coupling of mutant and wildtype genotypes with fluorescent labeling upon mitosis (Figure 1C-D). In a typical MADM experiment for the phenotypic characterization of the single cell loss of a candidate gene the homozygous mutant cells are routinely labelled in green fluorescent color (i.e. GFP^+^). For chromosomes with ‘forward’ MADM cassette configuration the mutant allele of a candidate gene must therefore be linked to the T-G MADM cassette. In such arrangement homozygous mutant cells will be labeled in green (GFP^+^) and wild-type cells in red (tdT^+^) (Figure 1C, Figure S1) upon a G2-X MADM event. Conversely, for chromosomes with ‘reverse’ MADM cassette configuration the mutant allele has to be linked to the G-T MADM cassette for the same genotype-labeling pattern (Figure 1D). In order to genetically link a mutant allele of a candidate gene to the corresponding chromosome containing the MADM cassette, meiotic recombination in germline can be utilized [e.g. (Hippenmeyer et al. 2010; Laukoter et al. 2020)] (Figure 1E-F). The probability for meiotic recombination that results in the linkage of the mutant allele with the MADM cassette can be estimated (Figure 1F) once the location (cM) of the mutant allele (genomic locus) has been determined by using for example Mouse Genome Informatics (MGI) database (www.informatics.jax.org).

Interestingly, homologous recombination frequencies in ES cells were relatively high for all selected loci (for some >50%), hinting at open chromatin structure which should be an advantage for prospective mitotic Cre-mediated interchromosomal recombination. Next, chimeric founder mice were generated by blastocyst injection. Homozygous *MADM*^*GT/GT*^ and *MADM*^*TG/TG*^ stock lines were established upon successful germline transmission of the respective MADM cassettes (Figure S3) by using specific genotyping primers (STAR Methods).

### Ubiquitous Labeling in all MADM Reporter Lines across Different Organs

Next we systematically analyzed the MADM labeling pattern upon Cre-mediated interchromosmal recombination in all MADM lines by introducing Cre driver lines (Figure S3E). First we crossed all *MADM*^*GT/GT*^ lines to mice that carry the Cre transgene within the X-linked *Hprt* (encoding hypoxanthine guanine phosphoribosyl transferase) genomic locus (Tang et al. 2002). *Hprt*^*Cre*^ is spatiotemporally ubiquitously and constitutively expressed (Tang et al. 2002). In female mice inactivation of the X chromosome results in mosaic *Cre* expression from the *Hprt* locus within tissues, and thus highly variable MADM labeling patterns (Hippenmeyer et al. 2013; Zong et al. 2005). We therefore analyzed male experimental MADM (*MADM*^*GT/TG*^*;Hprt*^*Cre*^/Y) animals for first pass comparative assessment of all MADM lines. We detected MADM labeling in all organs analyzed - including brain, spinal cord, eye, heart, lung, liver, kidney, thyme, and spleen (Figure 2A) - and in all MADM lines. The relative recombination frequency at least at this superficial qualitative level (see below for quantitative assessment) appeared to correlate in distinct selected organs across all 19 MADM (Figures 2–4 and S4–S5).

**Figure 2.**
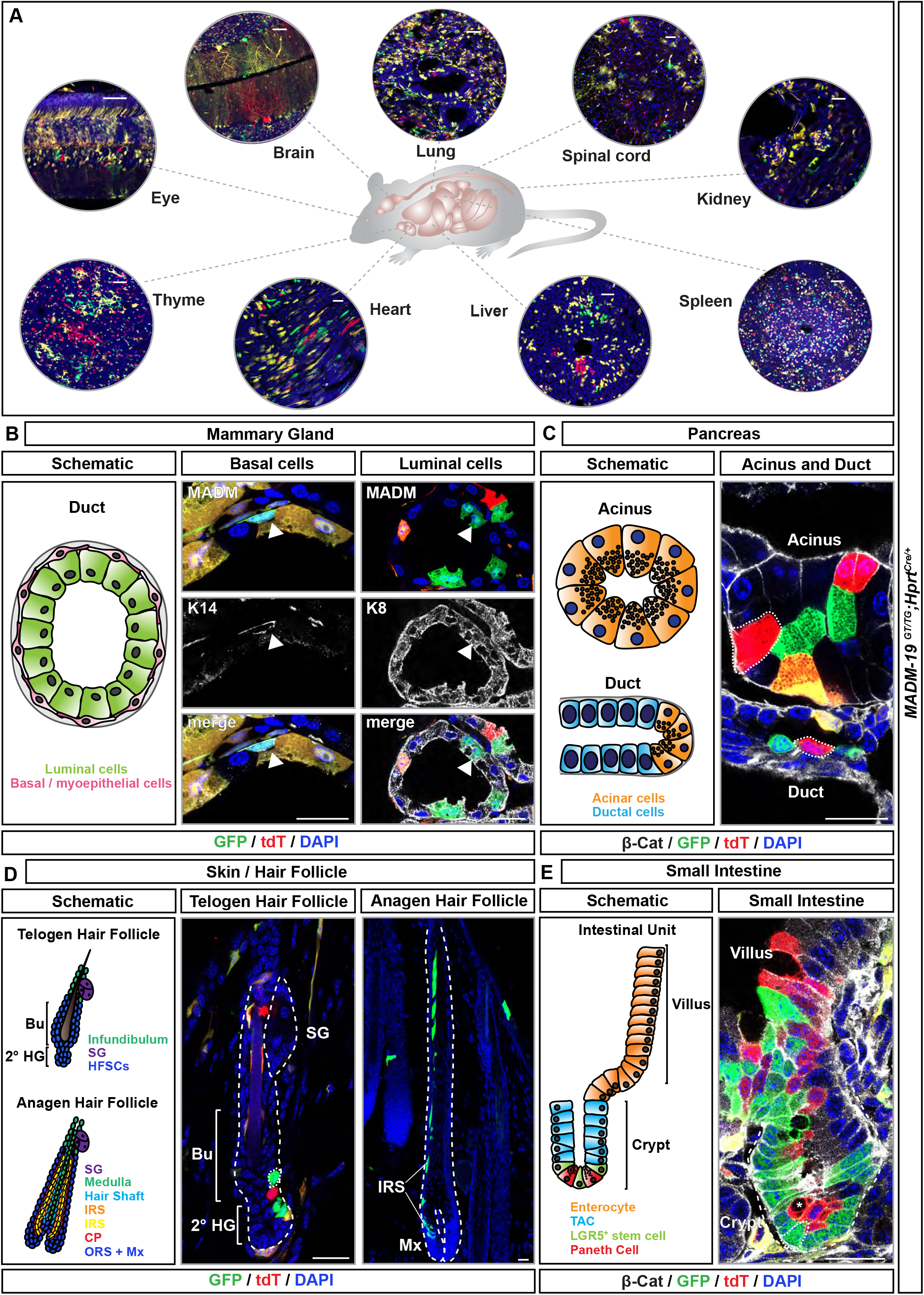
MADM Labeling Pattern in Different Organs and Stem Cell Niches. **(A)** Overview of MADM labeling (green, GFP; red, tdT; yellow, GFP/tdT) in *MADM-19*^*GT/TG*^ in combination with *Hprt*-Cre at P21. Diverse tissues and organs including eye, brain, lung, spinal cord, kidney, spleen, liver, heart, and thyme are illustrated. Scale bar: 50μm. **(B)** Schematic (left) and MADM labeling (middle/right; green, GFP; red, tdT; yellow, GFP/tdT) in mammary gland of lactating *MADM-19*^*GT/TG*^*;Hprt-Cre* female at four months of age. Basal/myoepithelial (middle) and luminal (right) cells are stained with antibodies against K14 and K8 (white), respectively. **(C)** Schematic (left) MADM labeling (right; green, GFP; red, tdT; yellow, GFP/tdT) in *MADM-19*^*GT/TG*^*;Hprt-Cre* pancreas, acinus and duct, at P21. Epithelial cells are visualized by antibody staining against β-Catenin (white, β-Cat). Acinar cells are identified by the presence of intracellular secretory granules. **(D)** Schematic (left) and MADM labeling (middle/right; green, GFP; red, tdT; yellow, GFP/tdT) in telogen (middle) and anagen (right) hair follicles in *MADM-19*^*GT/TG*^*;Hprt-Cre* at P21 (telogen) and P28 (anagen). Bu, bulge; 2° HG, secondary hair germ; SG, sebaceous gland; IRS, inner root sheath; CP, companion layer; ORS, outer root sheath; Mx, matrix. **(E)** Schematic (left) and MADM labeling (right; green, GFP; red, tdT; yellow, GFP/tdT) in small intestine in *MADM-19*^*GT/TG*^*;Hprt-Cre* at P21. Epithelial cells are visualized by antibody staining against β-Catenin (white, β-Cat). Asterisk marks a Paneth cell, identified by the presence of intracellular granules. TAC, transit-amplifying cell; LGR5, leucine-rich repeat-containing G-protein coupled receptor 5. Nuclei are stained using DAPI. Scale bar (B-E): 20μm.

### MADM Labeling in Clinically Relevant Adult Stem Cell Niches

MADM has been frequently utilized in the past to study lineage progression of progenitor stem cells in a variety of tissues. Furthermore, MADM has been used for the analysis of diseases with clonal origin. Most prominently, tumor formation and the delineation of cancer cell of origin at single cell level upon the introduction of mutations in tumor suppressor genes and/or loss of heterozygosity has been studied (Gonzalez et al. 2018; Liu et al. 2011; Muzumdar et al. 2016; Muzumdar et al. 2007; Yao et al. 2020). Given the enormous genome-wide potential inherent to the library of new MADM lines we next evaluated a number of stem cell niches with high clinical relevance. Since it is important to know the approximate scale of labeling for determining sample size in a MADM experiment, we chose two different new MADM models in combination with *Hprt*^*Cre*^ for these analyses: MADM-19 which shows relatively dense MADM-labeling, and MADM-4 which represents one of the sparsest MADM.

First, we focused on the mammary gland (Figure 2B), the site where breast cancer initiates. The mammary gland harbors two types of unipotent stem cell lineages, the K14^+^ myoepithelial (or basal) cells and the K8^+^ luminal cells (Van Keymeulen et al. 2011). Myoepithelial and luminal stem cell populations are derived from a multipotent progenitor during embryonic development (Wuidart et al. 2018), become unipotent at birth and can both give rise to mammary tumors upon transformation. For example, the frequently detected oncogenic *Pik3ca* (H1047R) mutation induces reprogramming of the unipotent progenitors to become multipotent cancer stem cells, thereby catalyzing the formation of heterogeneous, multi-lineage mammary tumors (Koren et al. 2015; Van Keymeulen et al. 2015). We evaluated MADM-labeling pattern in the postnatal mammary gland in adult lactating four month old female *MADM-19*^*GT/TG*^*;Hprt*^*Cre*/+^ (Figure 2B) and *MADM-4*^*GT/TG*^*;Hprt*^*Cre*/+^ (Figure S6A) mice and could readily detect GFP^+^ (green), tdT^+^ (red), and GFP^+^/tdT^+^ (yellow) cells in both K14^+^ basal and the K8^+^ luminal cells.

Next, we analyzed pancreatic epithelial cells which can be divided into secretory acinar cells and ductal epithelial cells. Although the tumor cell of origin for pancreatic cancer remains controversial, oncogenic drivers can trigger pancreatic ductal adenocarcinoma (PDAC) from both ductal and acinar cells (Ferreira et al. 2017; Lee et al. 2018). In both, *MADM-19*^*GT/TG*^*;Hprt*^*Cre/+*^ and *MADM-4*^*GT/TG*^*;Hprt*^*Cre/+*^ mice at P21 we noticed MADM-labeled cells in the acinus and duct within the pancreas (Figure 2C and Figure S6B).

Hair follicles are a prime stem cell model for the study of tissue regeneration but also for skin cancer including melanoma (Sun et al. 2019). Hair follicles are appendages of the epidermal lineage and undergo cycling rounds of stem cell activation in order to generate new hair (Fuchs and Nowak 2008). The stem cells are located in the secondary hair germ (2° HG) and lower part of the bulge (Bu) of a resting follicle (so called telogen follicle) (Figure 2D). They become activated, start to proliferate and expand the hair follicle deep down into the dermis. Progenitors located at the bottom of the activated follicle (so called anagen follicle) form the matrix, from where epithelial hair lineages are specified (Hsu et al. 2014). Such differentiated hair lineages comprise the companion layer (CP), distinct layers of inner root sheath (IRS), cuticle and cortex of the hair shaft (HS), as well as the innermost hair layer, the medulla (Me). Once hair regeneration is completed, the follicles undergo a destructive phase (so called catagen) and enter the quiescent resting phase again. In the skin of *MADM-19*^*GT/TG*^*;Hprt*^*Cre/+*^ and *MADM-4*^*GT/TG*^*;Hprt*^*Cre/+*^ mice we observed prominent MADM labeling in all compartments of the hair follicle and importantly in the hair follicle stem cells (Figure 2D and Figure S6C).

Lastly, we analyzed MADM labeling in the small intestine which represents another critical model for the study of stem cell mediated regeneration but also intestinal cancer (Barker et al. 2009). Intestinal stem cells replenishing the epithelium are LGR5^+^ and located in the crypt base (Barker et al. 2007). They are intermingled with secretory Paneth cells and divide constantly in order to rejuvenate the epithelial cell layer on the villus surface. Interestingly, LGR5^+^ stem cells mostly divide symmetrically and undergo neutral competition within the crypt, thus driving the crypt towards monoclonality (Snippert et al. 2010). In order to evaluate the potential for MADM-based lineage tracing, the study of loss of gene function, and analysis of stem cell behavior in the intestinal crypts we dissected the intestine of *MADM-19*^*GT/TG*^*;Hprt*^*Cre/+*^ and *MADM-4*^*GT/TG*^*;Hprt*^*Cre/+*^ mice at P21. We observed MADM-labeled cells in all compartments of the intestinal unit, including the villus and the crypt (Figure 2E and Figure S6D).

### Genomic Imprinting Phenotypes in Liver Cells with Uniparental Chromosome Disomy

MADM can create uniparental chromosome disomy (UPD, cells that carry two chromosomes from mother and maternally expressed imprinted genes are thus overexpressed whereas paternally expressed genes are not expressed and vice versa) (Figure 3A). This feature has been utilized to analyze imprinting phenotypes at single cell level that result from the imbalanced expression of imprinted genes in UPD (Hippenmeyer et al. 2013; Laukoter et al. 2020). Importantly, the analysis of candidate gene function, i.e. loss of function phenotypes, can be separated from UPD-mediated imprinting phenotypes by reverse MADM breeding schemes (Beattie et al. 2017; Hippenmeyer et al. 2013; Hippenmeyer et al. 2010; Joo et al. 2014; Laukoter et al. 2020). So far, prominent imprinting phenotypes have mostly been observed in liver where for instance cells with MADM-induced UPD of Chr. 7 exhibit paternal overgrowth (Hippenmeyer et al. 2013), in accordance with the kinship hypothesis that stipulates a major growth regulatory function of genomic imprinting (Haig 2004; Tucci et al. 2019). Since imprinted genes are located throughout the genome, we analyzed the livers in all 19 new MADMs in combination with *Hprt*-Cre (Figure 3B-3U) for potential additional imprinting phenotypes. We readily observed the paternal growth advantage of hepatocytes with paternal UPD of Chr. 7 (Figures 3H and 3V) but also noticed that cells with paternal UPD of Chr. 11 (Figures 3L and 3V) and Chr. 17 (Figure 3R and 3V) showed significant overrepresentation in comparison to cells with maternal UPD. Interestingly, the maternally expressed growth inhibitory imprinted genes *Grb10* and *Igf2r* are located on Chr. 11 and Chr. 17, respectively. Thus, while overexpression of growth promoting *Igf2* in UPD of Chr. 7 leads to paternal growth dominance (Hippenmeyer et al. 2013) the absence of growth antagonizing *Grb10* or *Igf2r* (Smith et al. 2006) may result in the growth advantage of cells with paternal UPD of Chr. 11 or Chr. 17 over cells with maternal UPD. We did not find significant UPD-mediated phenotypes in the livers of any other MADM besides MADM-7, MADM-11 and MADM-17 (Figure 3B-U), nor an indication for prominent imprinting phenotypes in any other organ analyzed at the current level of resolution.

**Figure 3.**
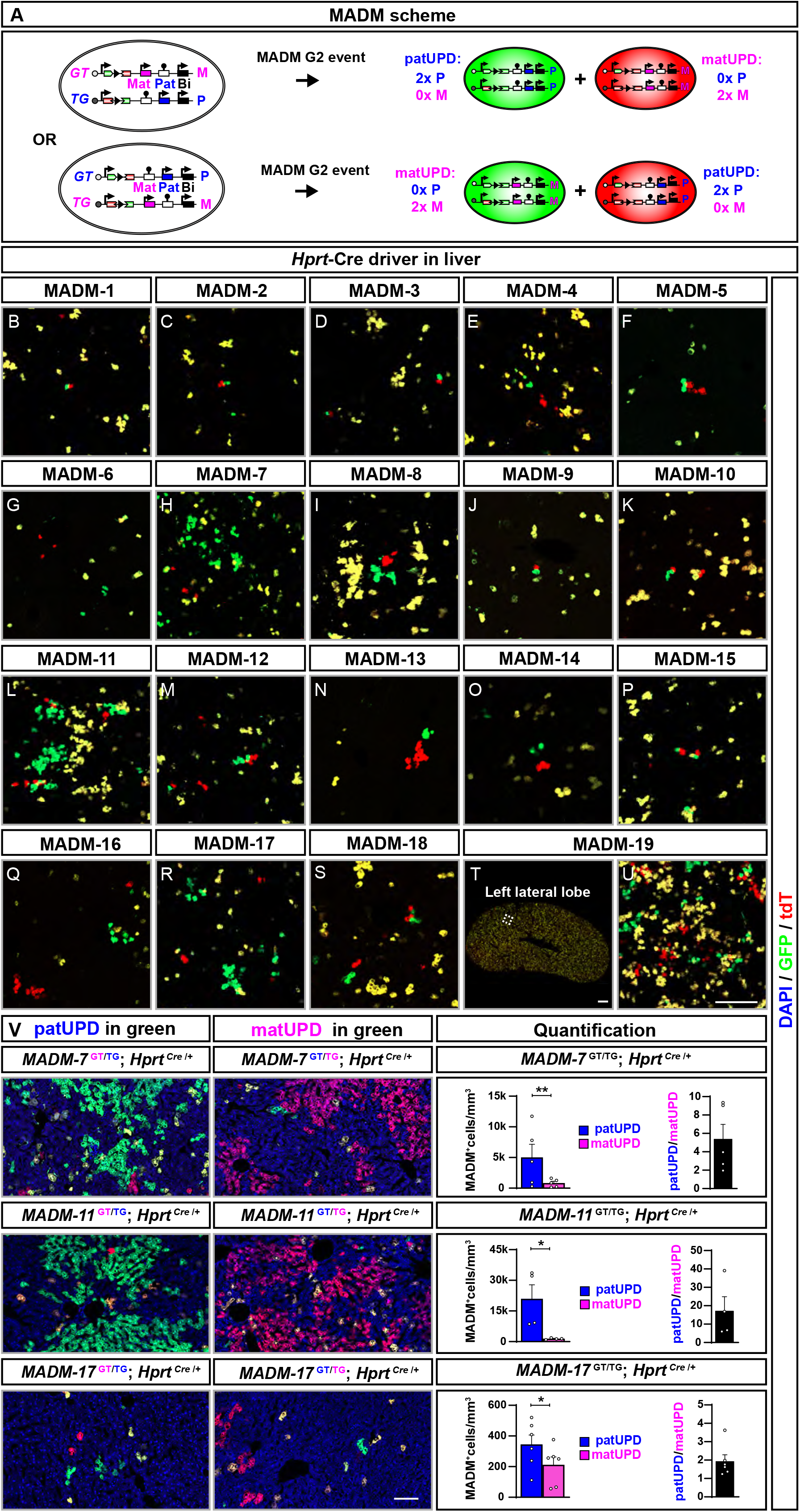
MADM-induced Uniparental Chromosome Disomy Results in Paternal Growth Dominance in Liver. **(A)** MADM scheme for imprinted genes. G2-X MADM events can generate differentially-labeled cells with near complete uniparental chromosome disomy [UPD, cells with two copies of either the maternal (matUPD) or the paternal (patUPD) chromosome], for details see also (Hippenmeyer et al. 2013; Laukoter et al. 2020). (Top) The GT MADM cassette is inherited from the mother (M, pink) and the TG MADM cassette from the father (P, blue). Consequently, green cells show patUPD (PP) and red cells matUPD (MM). In such scenario, imprinted maternally expressed genes are expressed at twice the normal dose and paternally expressed genes are not expressed in cells with matUPD (red). In contrast, paternally expressed genes are overexpressed by factor two and maternally expressed genes are not expressed in cells with patUPD (green). (Bottom) Reverse scheme where GT MADM cassette is inherited from father and TG MADM cassette inherited from mother. Here cells with matUPD are labelled in green and cells with patUPD in red fluorescent color. **(B-U)** Representative images of horizontal liver cryosections with MADM labeling (GFP, green; tdT, red) in MADM-1 (A) to MADM-19 (T-U) in combination with *Hprt*-Cre driver at P21. Higher resolution image (U) represents inset in (T) in left lateral lobe of liver in MADM-19. **(V)** (Top) Representative images (left, middle) of liver in *MADM-7*^*GT/TG*^*;Hprt-Cre* with green GFP^+^ patUPD and red tdT^+^ matUPD (left) or red tdT^+^ patUPD and green GFP^+^ matUPD (middle) at P21; and quantification (right) of absolute (#cells/mm^3^) and relative (PP/MM) numbers of MADM-labeled cells with UPD. (Middle) Representative images (left, middle) of liver in *MADM-11*^*GT/TG*^*;Hprt-Cre* with green GFP^+^ patUPD and red tdT^+^ matUPD (left) or red tdT^+^ patUPD and green GFP^+^ matUPD (middle) at P21; and quantification (right) of absolute (#cells/mm^3^) and relative (PP/MM) numbers of MADM-labeled cells with UPD. (Bottom) Representative images (left, middle) of liver in *MADM-17*^*GT/TG*^*;Hprt-Cre* with green GFP^+^ patUPD and red tdT^+^ matUPD (left) or red tdT^+^ patUPD and green GFP^+^ matUPD (middle) at P21; and quantification (right) of absolute (#cells/mm^2^) and relative (PP/MM) numbers of MADM-labeled cells with UPD. Nuclei are stained using DAPI. Scale bar: 200μm.

### Quantification of Recombination Efficiency of all MADM Chromosomes

The new MADM reporter mice appeared to exhibit distinct frequencies of Cre-mediated interchromosomal recombination albeit relative abundance of MADM labeling appears similar across different organs such as for instance brain (Figure 4 and S7), kidney (Figure S4) or spleen (Figure S5). To more rigorously and systematically determine recombination frequencies comparatively in all MADMs we quantified the absolute densities of MADM-labeled neurons in the cerebral cortex of P21 mice by using *Emx1*-Cre driver (Figures 5A-5B and S7). We first assessed MADM-labeling originating from G2-X events and quantified the numbers of green GFP^+^ and red tdT^+^ projection neurons per cubic millimeter (Figure 5A-5B). The relative numbers of red tdT^+^ versus green GFP^+^ projection neurons was not significantly different across MADM lines (Figures 5B and Table S1). Based on the density values we classified all the MADMs into three categories: 1) sparse with <25 cells/mm^3^; 2) intermediate with 25-100 cells/mm^3^ and 3) dense with >100 cells/mm^3^. The density of MADM-labeled cortical projection neurons in the densest MADM-11 was almost 2 orders of magnitude higher than in the sparsest MADM-4 (Figures 5B and Table S1). There was no correlation of the MADM recombination frequency with the size of the chromosome nor the density of genes located on a particular chromosome. Since all new MADM targeting loci have been selected by using the same criteria, the origin of the variability in recombination frequency across all MADMs is currently not clear. In mice, the pairing of homologous chromosomes in somatic cells is infrequent and under tight regulation, unlike in the fruit fly *Drosophila* (Apte and Meller 2012). Thus the spatial and/or dynamic organization of homologous chromosomes within the nucleus may result in distinct probabilities of Cre-mediated interchromosomal recombination in different MADM chromosomes. Regardless of the precise mechanism, all MADM reporters do work as predicted from the MADM principle (Figures 1 and S1) in the brain (Figures 4, 5A, 5B and S7) and all organs analyzed (Figures 2A, 3 and S4–S5). Importantly even the sparsest MADM-4 reliably permits functional genetic mosaic analysis of candidate genes (Hansen and Hippenmeyer; unpublished observation).

**Figure 4.**
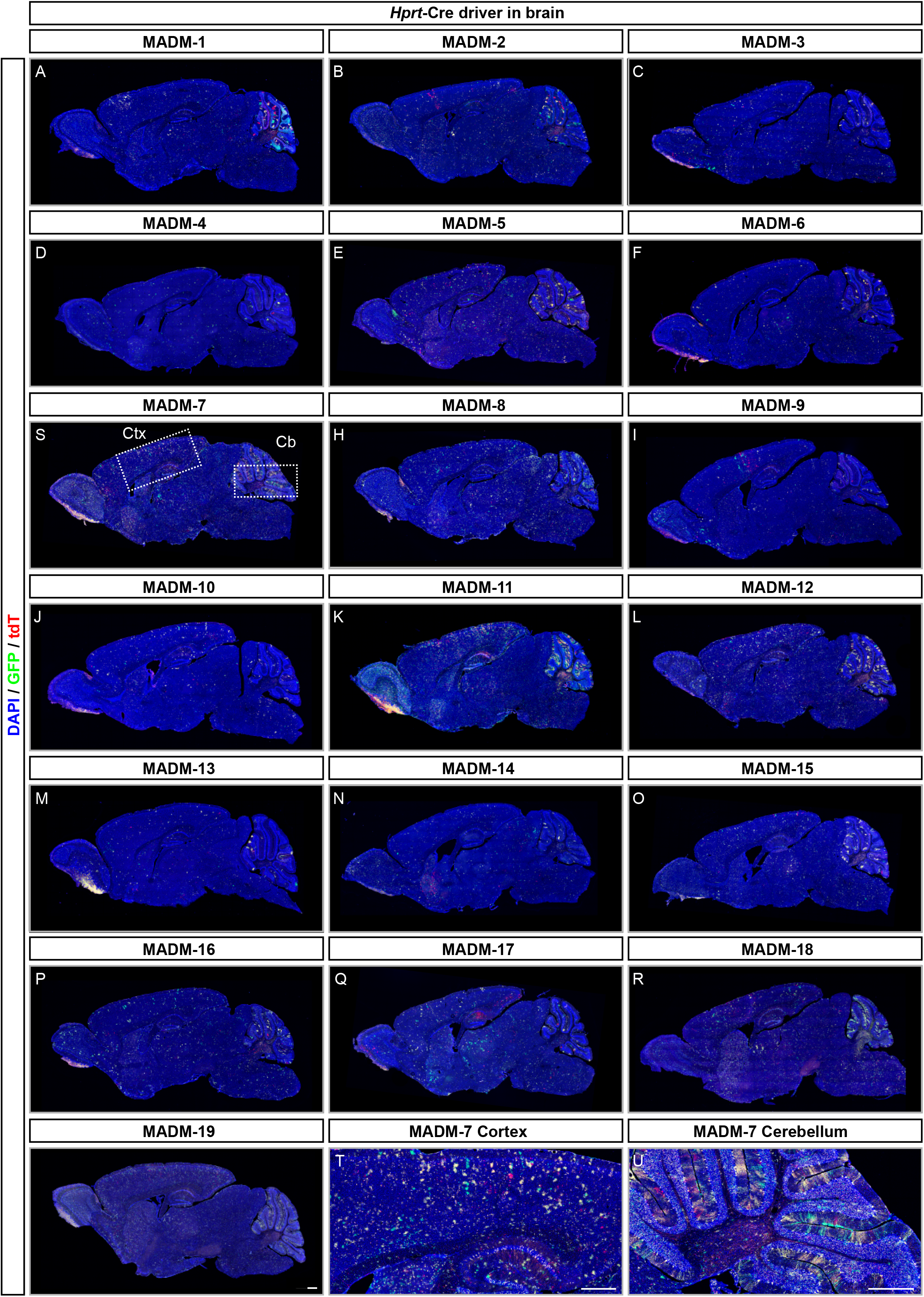
MADM Labeling Pattern in Brain across all 19 MADM Reporters. **(A-S)** Representative images of sagittal brain cryosections with MADM labeling (GFP, green; tdT, red) in MADM-1 (A) to MADM-19 (S) in combination with *Hprt*-Cre driver at P21. **(T-U)** Higher resolution images of cerebral cortex (T) and cerebellum (U) in MADM-7. Nuclei are stained by using DAPI (blue). Scale bar: 500μm.

**Figure 5.**
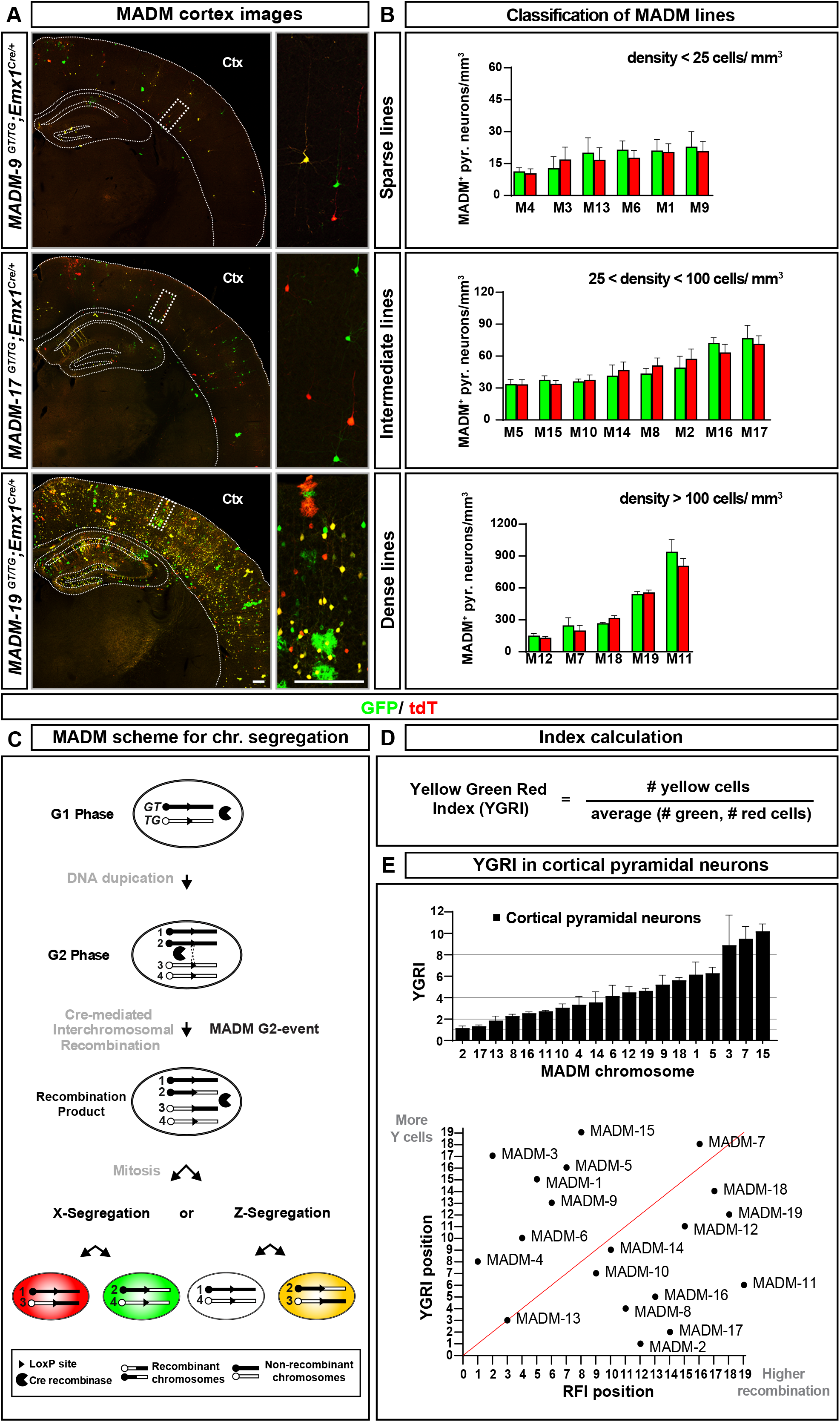
Mitotic Interchromosomal Recombination Efficiency and Sister Chromatid Segregation Patterns for all MADM Reporters in Cortical Projection Neurons. **(A)** Representative images of MADM labeling pattern (green, GFP; red, tdT) in cerebral cortex in three exemplary MADM lines in combination with *Emx1*-Cre driver at P21. Top, *MADM-9*^*GT/TG*^;*Emx1*^*Cre/+*^; middle, *MADM-17*^*GT/TG*^;*Emx1*^*Cre/+*^; bottom, *MADM-19*^*GT/TG*^;*Emx1*^*Cre/+*^. Scale bar:100 μm. **(B)** Classification of MADM lines. The 19 different MADM lines were classified by the efficiency of mitotic recombination reflected by the density of MADM labeling per mm^3^. Lines with less than 25 labeled cells per mm^3^ belong to the sparse category (top), lines with a density of labeled cells between 25 to 100 labeled cells per mm^3^ are in the intermediate category (middle), and lines with more than 100 labeled cells per mm^3^ belong to the dense category (bottom). **(C)** MADM principle for the monitoring of G2-X and G2-Z segregation patterns. Upon Cre-mediated interchromosomal recombination, at the LoxP site in the MADM cassettes, in G2 phase of the cell cycle recombinant chromosomes can either segregate together to the same daughter cell (G2-Z segregation; yellow, GFP/tdT and unlabeled cell) or each recombinant chromosome may segregate to distinct daughter cells (G2-X segregation; green, GFP and red tdT cells) upon mitosis. **(D)** Definition of YGR Index (YGRI). The YGRI is calculated from the number of yellow cells divided by the average of green and red cells to compensate for G2-Z events which leads to labeling of only one daughter cell (yellow) and an (invisible) unlabeled cell. Note that yellow cells emerging from G1/G_0_ events contribute to the total number of yellow cells. **(E)** YGR index in neuronal lineages. (Top) YGRI for cortical projection neurons in P21 neocortex of all 19 MADM reporter lines in combination with *Emx1*-Cre driver. Note that 1) YGRI varies from 1 to 10 but is never below 1; and 2) YGRI do not correlate with the sizes of the respective MADM chromosomes. (Bottom) YGRI ranking in correlation (red line) to recombination frequency index (RFI). Note that MADM chromosomes with a high recombination frequency do not necessarily present high YGRI and vice versa.

### MADM Reveals Chromosome-specific Biases in Mitotic Sister Chromatid Segregation Patterns

Previous *in vitro* studies have employed mitotic recombination in embryonic stem (ES) cells (and derived lineages) to monitor the randomness of sister chromatid segregation patterns upon mitosis (Armakolas and Klar 2006; Liu et al. 2002). Against common belief, initial results indicated that sister chromatids derived from a homologous pair of chromosomes did not segregate randomly to daughter cells. Instead, G2-X segregation (two recombinant chromosomes segregate away from each other), thus reflecting one particular pattern of sister chromatid segregation, prevailed in ES cells for Chr. 7 (Liu et al. 2002). Furthermore, ES cell-derived endoderm cell lines exhibited complete bias towards G2-X (Armakolas and Klar 2006). Conversely, ES cell-derived neuroectoderm cell lines never showed G2-X at all (Armakolas and Klar 2006). Although these results indicated that cell type may influence selective segregation of sister chromatids, such hypothesis is based on the analysis of only one chromosome and has not been examined in intact tissue *in vivo*. To this end we utilized the entire library of MADM-rendered homologous chromosomes to systematically trace sister chromatid segregation patterns of all 19 mouse autosomes in a number of somatic cell lineages *in vivo*.

We exploited the inimitable feature provided by the MADM principle (Figure 5C and S1) – the differential fluorescent labeling of pairs of nascent sister cells upon mitosis which is dependent on how recombinant chromosomes segregate during cell division. G2-X segregation of recombinant MADM chromosomes can be unambiguously identified (by the presence of red and green cells). However, G2-Z segregation, producing yellow cells, cannot be identified without ambiguity because G1 and/or postmitotic G_0_ events also result in yellow cells (Zong et al. 2005) (Figure 5C and S1). Therefore we capitalized upon the power of unequivocal G2-X identification - but also taking into consideration the caveat of yellow cells potentially reflecting a mix of G2-Z and G1/G_0_ - and defined ‘yellow-green-red-index’ (YGRI) as a proxy for sister chromatid segregation patterns (Figure 5D).

First, we systematically determined the YGRI of pyramidal projection neurons in the P21 neocortex for all 19 MADM reporters in combination with *Emx1*-Cre driver (expressed in cortical progenitor cells and thereby limiting G_0_ events) (Figure 5E and Table S1). Contrary to the prediction and expectation based on cell culture data [no G2-X in neuroectodermal lineage (Armakolas and Klar 2006)] we always observed G2-X events. Interestingly, the YGRI values ranged from ~1 for MADM-2 and MADM-17 to ~10 for MADM-15 (Figure 5E, top). The values of the YGRI did not correlate with the sizes of the respective MADM chromosomes. Next we compared the values of the YGRI with the absolute recombination frequencies (RFI, recombination frequency index), i.e. density of G2-X MADM labeling as indicated in Figure 5B. In the ranking plot where axes indicate YGRI versus RFI, there was no apparent correlation (Figure 5E, bottom) of YGRI with RFI. In other words, MADM chromosomes that showed a high recombination frequency did not necessarily present with a high YGRI and vice versa (Figure 5E, bottom). In summary, we detected highly distinct YGRI for different MADM chromosomes, suggesting distinct sister chromatid segregation patterns in cortical *Emx1*^+^ projection neuron lineage.

### Chromosome-specific Biases of Sister Chromatid Segregation Differ in Distinct Cell Types

To determine the influence of cell type on biased, chromosome-specific, sister chromatid segregation patterns we first analyzed cortical astrocytes and hippocampal CA1 pyramidal cells, both of which are derived from *Emx1*^+^ progenitor cells. The YGRI for cortical astrocytes was markedly different from the YGRI for cortical projection neurons or hippocampal CA1 pyramidal cells for a representative set of ten MADM chromosomes analyzed (Figure 6A). Interestingly, YGRI of all 10 chromosomes in astrocytes were rather constant and low, indicating a uniformly high relative frequency for G2-X events in astrocyte progenitors. Next we introduced *Nestin*-Cre into the same ten MADM reporters to label neural lineages beyond forebrain projection neurons and astrocytes. We focused on cerebellar Purkinje cells which can be unambiguously identified by their characteristic morphology and determined the YGRI for these cells. Strikingly, the YGRI for Purkinje cells was also markedly different in most MADMs when compared to the YGRIs for cortical projection neurons and astrocytes, and hippocampal CA1 pyramidal cells (Figure 6A). Moreover, chromosomes that have high YGRI in one cell type would not necessarily show high YGRI for a different cell type.

**Figure 6.**
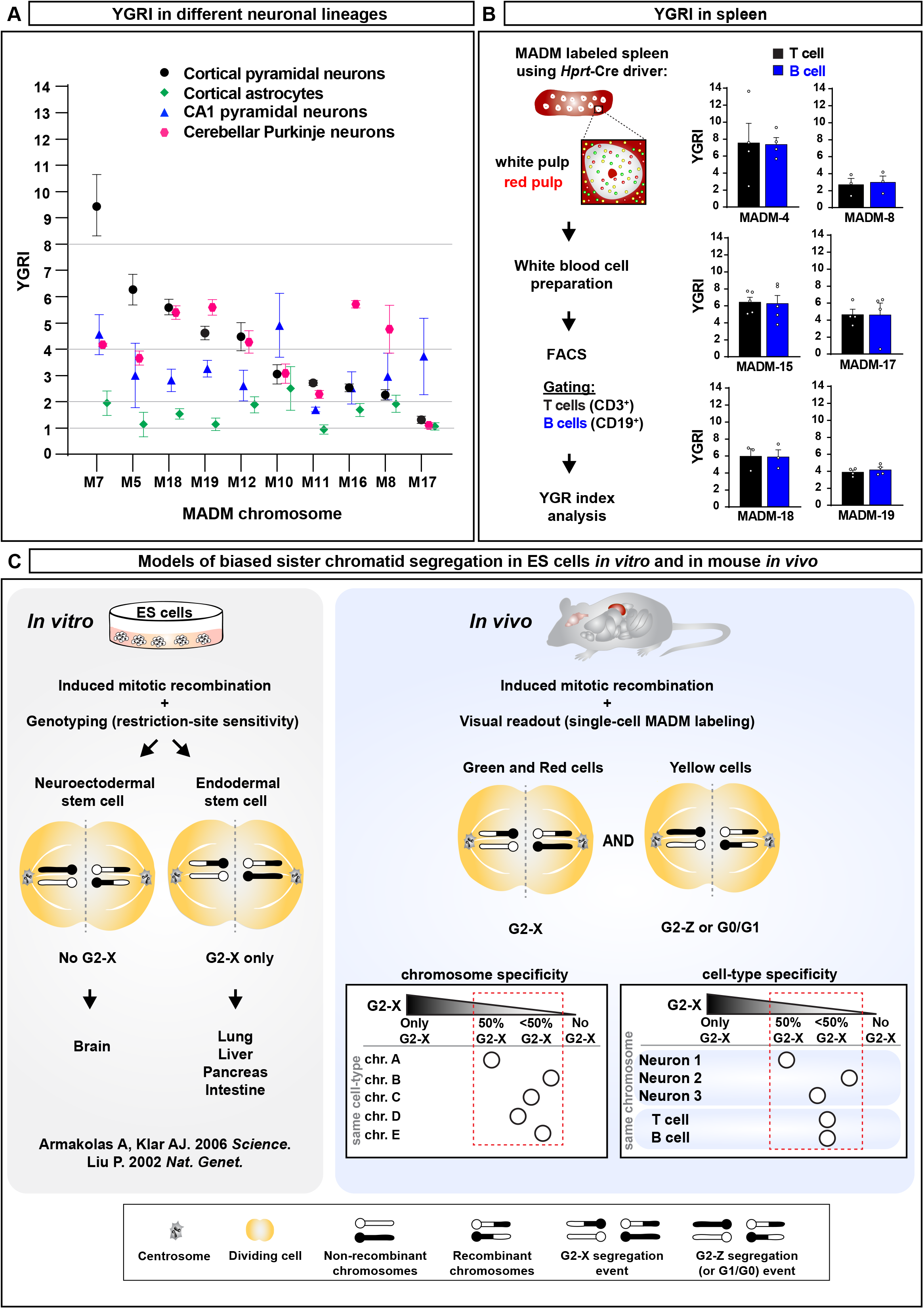
Sister Chromatid Segregation Patterns Differ in Distinct Somatic Cell Lineages. **(A)** YGRI for selected MADM reporters in different neuronal lineages. YGRI of cortical astrocytes and hippocampal CA1 pyramidal cells derived from *Emx1*^+^ progenitors and cerebellar Purkinje cells derived from *Nestin*^+^ progenitors significantly differs from the YGRI of cortical pyramidal neurons for most MADM chromosomes analyzed. **(B)** (*Left*): White blood cell preparations from spleen in diverse MADM reporters in combination with *Hprt*-Cre at P21 were subjected to FACS. The number of green GFP^+^, red tdT^+^, and yellow GFP^+^/tdT^+^ CD3^+^ T cells (black) and CD19^+^ B cells (blue) were quantified. (*Right*) YGRI for six different MADM chromosomes including sparse (MADM-4), intermediate (MADM-8, MADM-15, MADM-17) and dense (MADM-18, MADM-19) lines. While the distinct MADM recombinant chromosomes displayed different YGRI values, the YGRI for T cells and B cells was not significantly different for all MADM chromosomes analyzed. Two-tailed Student’s t-test, p_M4_=7.4E-01, p_M8_=7.9E-01, p_M15_=7.7E-01, p_M17_=6.3E-01, p_M18_=9.8E-01, p_M19_=5.0E-01. **(C)** Models of biased sister chromatid segregation patterns in ES cells *in vitro* and in mouse *in vivo*. (Left) Previous studies (Armakolas and Klar 2006; Liu et al. 2002) employing mitotic recombination and in combination with restriction-site sensitivity for genotyping in ES cell cultures reported that in ES cell-derived neuroectodermal lineages no G2-X (recombinant chromosomes segregate away from each other during cell division) events could be observed. In contrast, lineages derived from endodermal stem cells showed exclusively G2-X segregation patterns. Based on these findings it could be anticipated that in MADM there would be no red and green cells in neural lineages (e.g. in the brain) which was not the case for all MADM chromosomes. (Right) *In vivo* analysis of the prevalence of G2-X events (red and green cells) in comparison with total number of yellow cells (G2-Z, G1 and G_0_ events) for all MADM chromosomes and in several somatic cell lineages revealed significant bias in recombinant chromosome, and thus sister chromatid segregation patterns. The segregation bias showed marked chromosome specificity: was distinct for different chromosomes in the same cell type in both brain and hematopoietic systems. The segregation bias appears also to be affected by cell type: the level of bias was distinct for the same chromosome in different cell types.

Finally, we assessed sister chromatid segregation patterns for a non-neural somatic cell type. We focused on T-cells (CD3^+^) and B-cells (CD19^+^) within the hematopoietic lineage, isolated MADM-labeled cells by FACS, and determined the YGRI for six different MADM chromosomes (Figure 6B). Interestingly, while the distinct MADM chromosomes showed different YGRI values, the YGRI for T-cells in comparison to B-cells was not significantly different for all six chromosomes analyzed. No significant correlation could be established when the YGRI of T/B-cells was compared to the YGRI of the neural lineages. Altogether, these data indicate that the highly biased and chromosome-specific sister chromatid segregation patterns are further affected by cell type in somatic cell lineages *in vivo*.

## DISCUSSION

The analysis of gene function in multicellular systems *in vivo* requires quantitative and high-resolution experimental tools and assays to analyze the cellular phenotype. MADM technology offers a genetic approach in mice to visualize and concomitantly manipulate genetically defined cells at clonal level and with single cell resolution. Here we expanded the MADM technology in order to enable the genetic dissection of cell-autonomous gene function of most genes (>96%) across the entire mouse genome. While functional genetic mosaic analysis clearly represents the most salient utility of MADM (Figure 1, S1 and Table 1) here we also extended the application spectrum, and utilized MADM as a proxy to trace randomness of mitotic sister chromatid segregation patterns for the first time *in vivo*. We first discuss these findings in a more general context before we elaborate on the overarching potential of the genome-wide MADM resource for future genetic mosaic analysis.

### Non-random Mitotic Sister Chromatid Segregation in Mouse *in vivo*

Asymmetric stem cell division requires the non-equivalent distribution of cell-fate determinants including proteins, mRNA or intracellular organelles (Gonczy 2008; Knoblich 2008; Taverna et al. 2014). Recently, an intriguing model has been postulated whereby asymmetric cell division might be also promoted by differentiation of sister chromatids by epigenetic means, followed by selective segregation of ‘unequal’ sister chromatids to daughter cells (Armakolas et al. 2010; Bell 2005; Yamashita 2013). However, experimental evidence supporting such a model in mice was thus far obtained solely from *in vitro* studies in ES cells and derived lineages, and only for one chromosome (Chr. 7; Figure 6C, left) (Armakolas and Klar 2006; Liu et al. 2002). In our study we systematically traced sister chromatid segregation patterns of the entire set of mouse autosomes, for the first time to our knowledge in a mammalian species *in vivo*. By capitalizing on the distinct fluorescent labeling of daughter cells upon MADM-based mitotic recombination we could unambiguously identify G2-X segregation (recombinant MADM chromosomes segregate away from each other), reflecting one particular pattern of sister chromatid segregation. Interestingly, we observed that the prevalence of G2-X events, reflected in the value of YGRI, in the same cell type (cortical projection neurons) and by using identical *Emx1*-Cre driver vastly differed, up to one order of magnitude for different chromosomes. Thus sister chromatid segregation is highly biased in a chromosome-specific manner in mitotic cortical *Emx1*^+^ progenitors. Furthermore, the rank orders of YGRI for each chromosome in different cell types were not the same, suggesting that the bias of sister chromatid segregation patterns results from a complex combination of chromosome and cell type specific mechanisms (Figure 6C, right).

Previous studies found that cultured ES cell clones that were differentiated into neuroectoderm lineage never showed G2-X segregation (Armakolas and Klar 2006; Liu et al. 2002). These findings are in stark contrast to our *in vivo* results demonstrating for all 19 mouse autosomes a substantial amount of G2-X segregation, and in at least four distinct neural cell lineages. While we cannot fully explain the cause of the differences in results obtained in cell culture or *in vivo*, respectively, systemic and/or tissue-wide acting mechanisms could be involved (Knouse et al. 2018). For our MADM-based analysis we used *Emx1*- and *Nestin*-Cre drivers which are mostly active in dividing neural stem cells and turned off in postmitotic cells. Contribution of G_0_ recombination is thus expected to be minimal. Still, all YGRIs in neural lineages were ≥1 (with some up to an order of magnitude higher) indicating increasing rates of G2-Z segregation. However, a certain rate of G1 recombination (also producing yellow cells that increase the YGRI) besides G2-Z segregation may add to the overall YGRI. Although G1 recombination events did not occur in cultured ES cells (Armakolas and Klar 2006; Liu et al. 2002), we cannot currently exclude that interchromosomal recombination efficiency could be distinct in G1 versus G2 phases of the cell cycle for different cell types *in vivo*. However, for any given cell division cycle, the relative recombination events in G1 versus G2 should be the same; thus, different YGRIs for different chromosomes must reflect chromosome-specific sister chromatid segregation patterns. It will be interesting in future to test whether the bias of sister chromatid segregation could be influenced by the location of the genomic recombination loci for particular chromosomes.

The putative underlying molecular mechanisms of biased sister chromatid segregation have been previously explored using *in vitro* assays. As such, the left-right dynein (LRD) protein was implicated in the selective sister chromatid segregation process (Armakolas and Klar 2007). These results are insofar intriguing since mutation of the gene (*Dnah11*) encoding LRD causes randomization of left-right laterality mice (half of the animals develop with mirror-imaged visceral organs). It will thus be interesting in future experiments to systematically assess the role of *Dnah11* in biased sister chromatid segregation *in vivo* across distinct somatic cell lineages by using the MADM approach.

The phenomenon of biased sister chromatid segregation appears to be evolutionarily conserved (Beumer et al. 1998; Pimpinelli and Ripoll 1986). Interestingly, in asymmetrically dividing male germline stem cells in *Drosophila*, sister chromatids of X and Y, but not autosomes are segregated non-randomly (Yadlapalli and Yamashita 2013). In such context, SUN-KASH proteins, proposed to anchoring of sister chromatids to centrosome, seem to be involved, besides regulators of DNA methylation (Yadlapalli and Yamashita 2013). While the underlying molecular mechanisms may or not be conserved, it will be intriguing to assess the physiological function in future studies, and experimentally approach the hypothesis postulating that biased sister chromatid segregation could be a mechanism to instruct cell fate of nascent daughter cells during asymmetric stem cell division (Armakolas et al. 2010; Bell 2005; Yadlapalli and Yamashita 2013). Since MADM enables both clonal lineage tracing with concurrent genetic manipulation, such approach promises high potential to systematically address the physiological role of biased sister chromatid segregation in future.

### Genome-wide MADM Mice Library for Unique Single-Cell Genetic Mosaic Analysis

#### Genetic dissection of cell-autonomous gene function and system-wide effects

The MADM technology enables a variety of genetic *in vivo* paradigms to study a broad spectrum of cell and developmental processes (Hippenmeyer 2013; Hippenmeyer et al. 2013; Luo 2007; Muzumdar et al. 2007; Zong et al. 2005). Here we expanded the most salient property of MADM, functional analysis of candidate gene function at single cell level, for the study of nearly all genes encoded in the mouse genome. One exclusive application of the MADM system is the feature enabling the genetic dissection of the relative contributions of cell-autonomous and extrinsic systemic and/or tissue-wide components to the overall cellular phenotype upon the loss of candidate gene function. Importantly, the single cell phenotype in conditional or full knockout mutants reflects a combination of both cell-autonomous gene function and environment-derived cues which may remedy or exacerbate any observed phenotype. In fact, recent genetic mosaic studies indicate that non-cell-autonomous mechanisms fundamentally impact developmental processes. For instance, MADM-based analysis of the genes encoding components of the cytoplasmic LIS1/NDEL complex in comparison with conditional knockout revealed highly specific sequential cell-autonomous functions for *Lis1* and *Ndel1* genes. In contrast, non-cell-autonomous effects resulting from whole tissue *Lis1*/*Ndel1* ablation appear to impact neuronal migration much more profoundly and throughout the entire process of cortex development (Hippenmeyer 2014). Furthermore, while *Lgl1*, encoding a regulator of cellular polarity, cell-autonomously controls cortical glia production, its function is also required at the global tissue level during embryonic neurogenesis and to prevent emergence of periventricular heterotopia (a.k.a. double cortex syndrome in human) (Beattie et al. 2017). Thus, insights at single cell resolution as obtained from MADM-based approaches in combination with systematic candidate gene interrogation (Beattie et al. 2017; Laukoter et al. 2020) likely will have implications for our general understanding of neurodevelopmental disorders such as Lissencephaly and related cortical malformations including microcephaly and hemimegancephaly among others (Buchsbaum and Cappello 2019; D’Gama and Walsh 2018; Jayaraman et al. 2018; Pinson et al. 2019; Subramanian et al. 2019).

#### Single cell analysis of imprinting phenotypes in uniparental chromosome disomy

A unique MADM application includes the property to generate cells with uniparental chromosome disomy (UPD) and thus enable the study of imprinting phenotypes at single cell level (Hippenmeyer et al. 2013; Laukoter et al. 2020). In fact, technical limitations so far only allowed the investigation of UPD at the whole animal level but lacked the resolution to obtain insights at the cellular level. Another major drawback in the analysis of UPD in whole animals is reflected in the key importance of many imprinted genes in nutrient transfer during pregnancy (Barlow and Bartolomei 2014). Thus the phenotypic interpretation of UPD at the individual cell level is confounded by putative whole animal wide systemic effects. MADM technology provides a solution and is to date the sole technology that can produce UPD sparsely in genetic mosaic animals and within genetically-defined cell populations (Hippenmeyer et al. 2013; Laukoter et al. 2020). It will be revealing in future studies to systematically probe the cell-autonomous consequences of UPD at single cell level and without inducing global changes in imprinted gene expression affecting the whole animal. The library of all nineteen MADM reporters will in principle enable the systematic analysis of UPD-associated cellular phenotypes in any organ, tissue and cell-type in the mouse, provided the availability of appropriate tissue-and/or cell type specific Cre drivers. In a broader context, since UPD in human is associated with a variety of diseases (Buiting et al. 2016; Feinberg 2007; Tuna et al. 2009; Yamazawa et al. 2010) MADM-based analysis will also contribute to our general understanding of the underlying etiology of imprinting disorders at single cell level.

#### Analysis of cellular competition at single cell level in health and disease

MADM can be exploited for the study of cellular competition in developmental context. For instance, when the TrkC Neurotrophin receptor is removed sparsely with MADM from just a few individual Purkinje cells in the cerebellum, their dendrites have fewer and shorter branches. In contrast, when TrkC is ablated from all Purkinje cells, the dendrite trees look normal again. Thus a competitive mechanism could be involved whereby the shape of the dendrite tree depends on relative differences in Neurotrophin/TrkC signaling between Purkinje cell neighbors (Joo et al. 2014). Cell competition has not only been implicated in cell morphogenesis but extensively studied in a variety of contexts. Cell competition is particularly critical for overall tissue homeostasis during growth and regeneration but also for cell mixing and tissue invasion in cancer (Bras-Pereira and Moreno 2018; Ellis et al. 2019; Madan et al. 2018; Merino et al. 2016). With the availability of MADM for all mouse autosomes the phenomenon of cell competition can be studied holistically and for virtually any candidate gene function associated with it in diverse biological contexts in health and disease.

#### Expansion to other species and future development of the MADM technology

While MADM technology currently is available only in mice future expansion of the system to other species by for instance CRISPR/Cas9-enabled transgenesis can be anticipated. The range of species suited for MADM depends however on a number of factors such as the ease of breeding and generation time. The study of genetic mosaicism in human context is another prospective application for the MADM system. The MADM cassettes could be inserted in human embryonic stem cell lines which then could be used for differentiation, generation and the study of cellular organoids in broad applications. With the implementation of numerous protocols for next generation sequencing, gene expression can be monitored in MADM-labeled cells in bulk and also at single cell level [Pauler and Hippenmeyer, unpublished observation, (Laukoter et al. 2020)]. Thus MADM-based cell lineage tracing in combination with single cell RNA-sequencing and *in silico* modeling holds the potential to holistically reconstruct cell lineages across all organs in the mouse. Lastly, the MADM reporter genes can be adapted for the monitoring of physiological processes by for instance intrinsic calcium imaging and/or optogenetic manipulation. Altogether, the whole mouse genome MADM resource presented in this study likely will catalyze the genetic dissection of the cellular and molecular mechanisms with single cell resolution across a broad spectrum of biological questions in health and disease.

## ACKNOWLEDGMENTS

We thank the Bioimaging-, Life Science- and Pre-Clinical Facilities at IST Austria; MP. Postiglione, C. Simbriger, K. Valskova, C. Schwayer, T. Hussain, M. Pieber, and Victoria Wimmer for initial experiments, technical support and/or assistance; M. Sixt, and all members of the Hippenmeyer lab for discussion; and W. Zhong for sharing *Nestin-Cre* mice. This work was supported by National Institutes of Health grants (R01-NS050580 to L.L. and F32MH096361 to L.A.S.). L.L. is an investigator of HHMI. N.A. received support from FWF Firnberg-Programm (T 1031). A.H. is a recipient of a DOC Fellowship (24812) of the Austrian Academy of Sciences. This work also received support from IST Austria institutional funds; FWF SFB F78 to S.H.; the People Programme (Marie Curie Actions) of the European Union’s Seventh Framework Programme (FP7/2007-2013) under REA grant agreement No 618444 to S.H., and the European Research Council (ERC) under the European Union’s Horizon 2020 research and innovation programme (grant agreement No 725780 LinPro) to S.H.

## Author Contributions

S.H. and L.L. conceived the research. S.H., L.L., L.A.S., T.R., N.A. and X.C designed all experiments and interpreted the data. X.C., A.D., N.A., A.H.H., J.S., L.A., T.B., T.R., L.A.S., A.H., R.J. performed all the experiments. S.H. wrote the manuscript with inputs from L.L and X.C. All authors edited and proofread the manuscript.

## SUPPLEMENTAL INFORMATION

### Supplemental Figures

**Figure S1.**
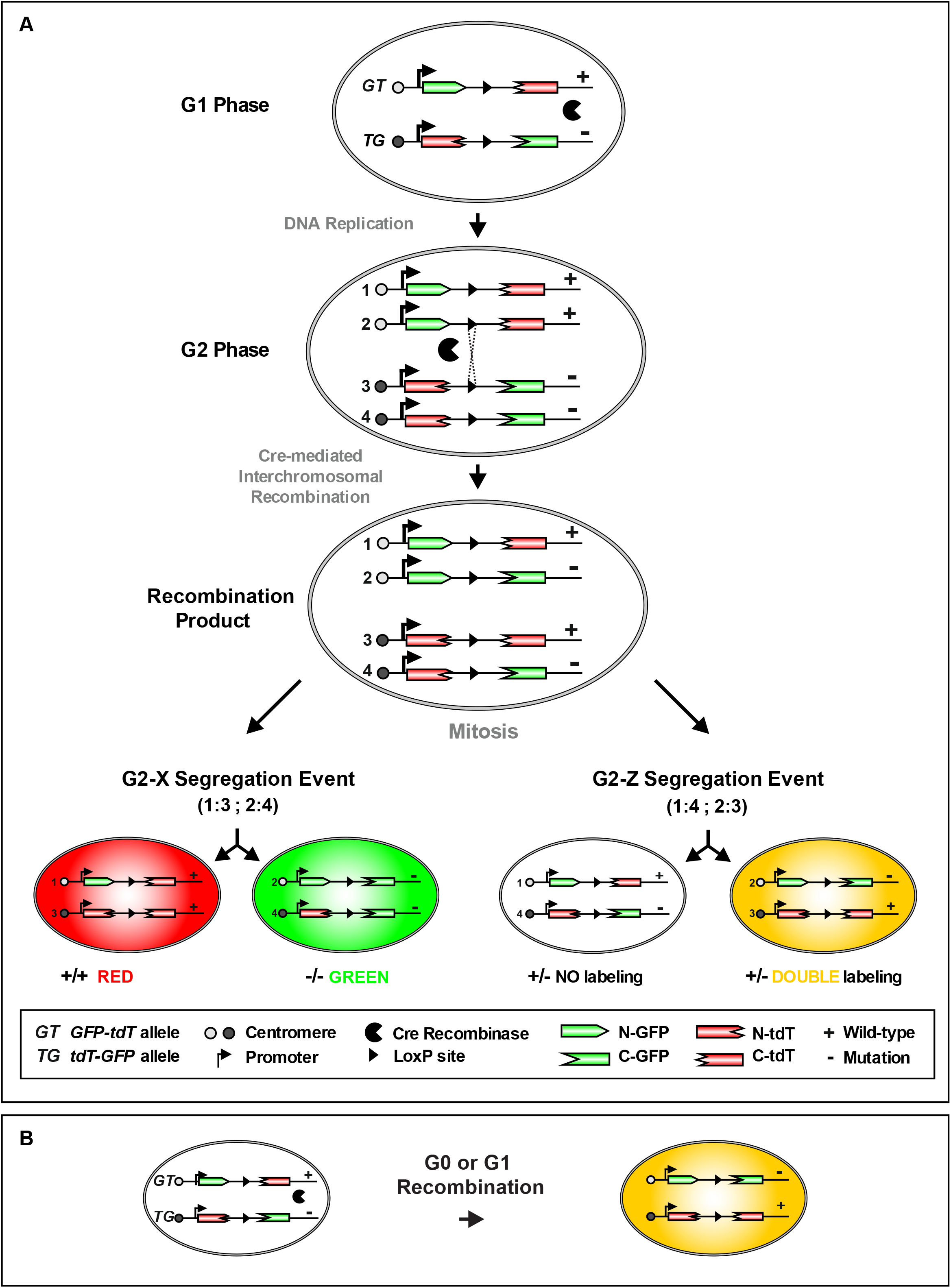
MADM Principle. **(A)** For MADM, two reciprocally chimeric marker genes are targeted to identical loci on homologous chromosomes. The chimeric marker genes (*GT* and *TG* alleles) consist of partial coding sequences for green (eGFP[G]) and red (tdT[T]) fluorescent proteins separated by an intron containing the loxP site. Following Cre recombinase-mediated interchromosomal recombination during mitosis, functional green and red fluorescent proteins are reconstituted resulting in two daughter cells each expressing one of the two fluorescent proteins upon G2-X events (recombination in G2 of the cell cycle followed by X segregation). Introduction of a mutation distal to one MADM cassette allows the generation of genetic mosaics with single cell resolution, with wild-type daughter cells, labeled with one color (e.g. red) and homozygous mutant siblings with the other (e.g. green) in an unlabeled heterozygous environment. G2-Z segregation results in yellow cell labeling with no genotype alteration. **(B)** G1 and G_0_ events also result in yellow cell labeling without change in genotype. Reproduced and adapted with permission from (Hippenmeyer et al. 2010).

**Figure S2.**
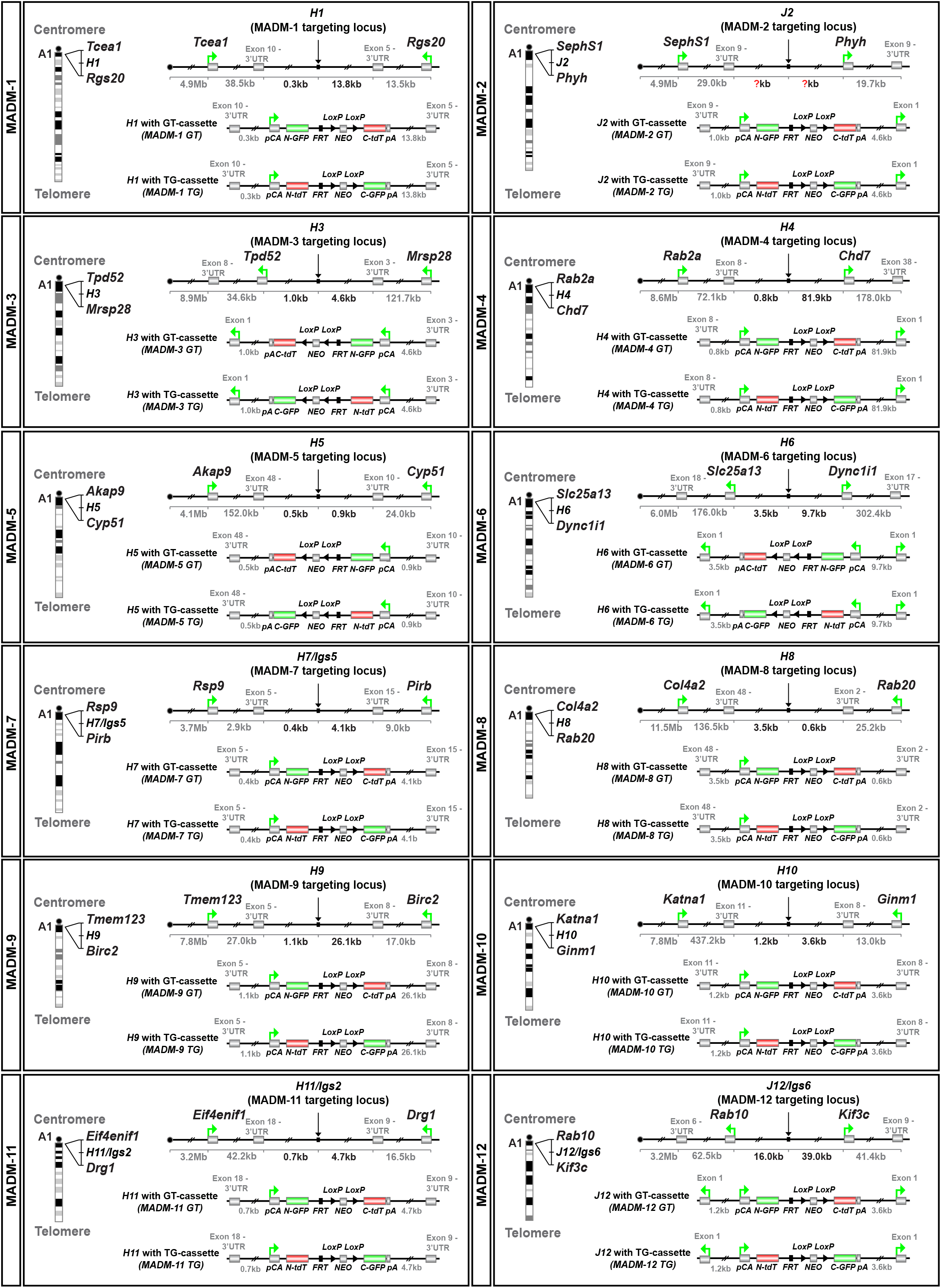
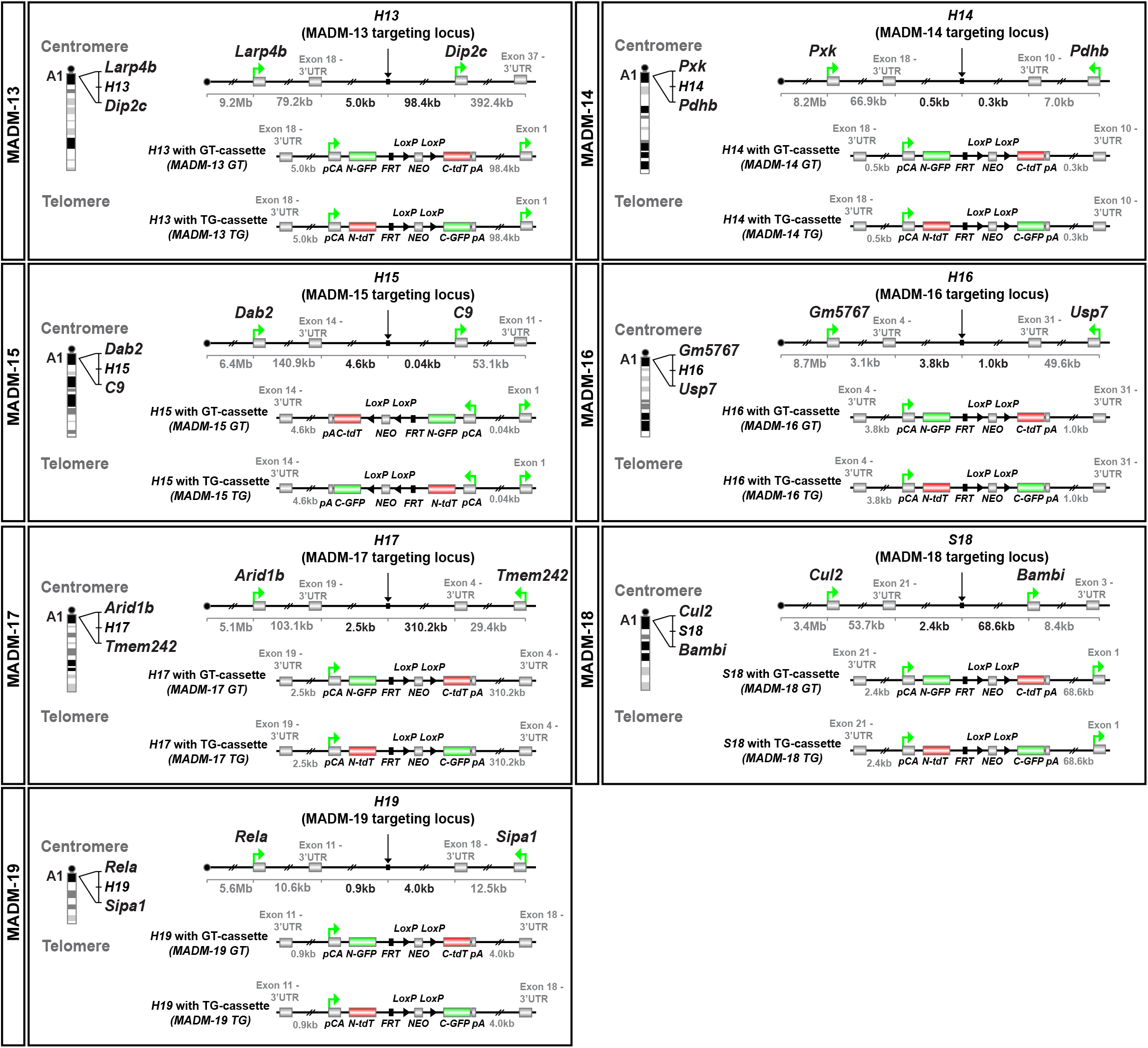
Genomic Targeting of GT and TG MADM Cassettes. For each MADM targeting site: in the panel on the left the schematic indicates the corresponding chromosome, insertion site of the MADM cassettes, and flanking genes. In each panel on the right a schematic illustrates the MADM cassette insertion site with genomic locus information (top) and the targeted locus with the G-T (middle) and T-G (bottom) MADM cassettes, respectively.

**Figure S3.**
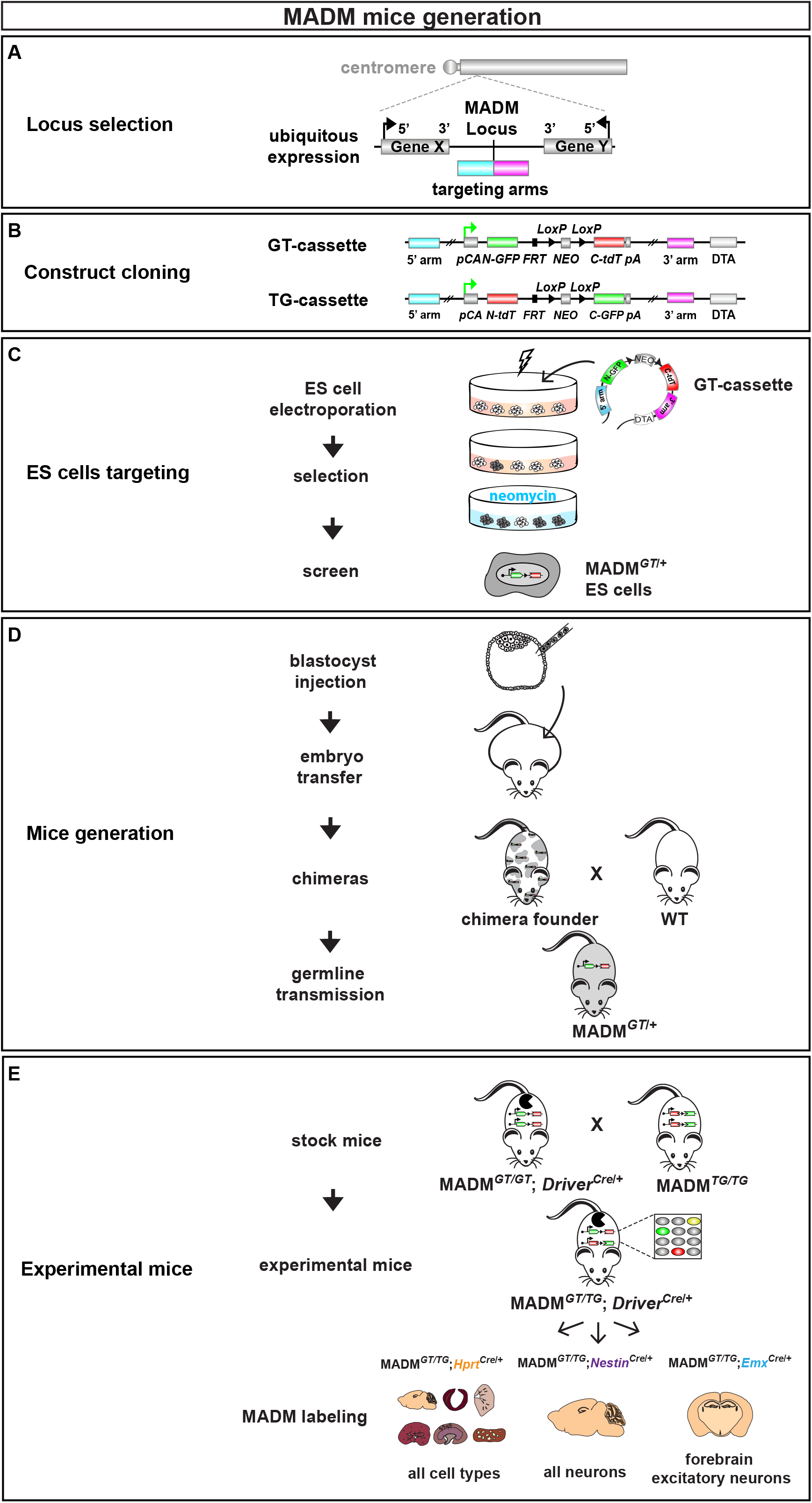
Workflow for the Generation of MADM Transgenic Mice. **(A)** Identification and selection of genomic targeting locus. The 3 key criteria for a suitable locus were 1) closest possible proximity to centromere to maximize the number of genes located distally to the MADM cassettes 2) intergenic location to minimize the probability of disrupting endogenous function and 3) permit spatially and temporally ubiquitous and biallelic expression of reconstituted GFP and tdT markers based on EST (expression sequence tag) data from UCSC Genome Browser. **(B)** Cloning of targeting vectors. Both GT-MADM cassette and TG-MADM cassette sequences were cloned separately into plasmids containing the 5’ and 3’ targeting arms. **(C)** ES cell targeting by electroporation. Targeting constructs for both GT-MADM cassettes and TG-MADM cassettes containing a diphtheria toxin A fragment and a positive selection marker (neomycin resistance cassette) were electroporated separately into ES cells. After selection, ES clones were picked and tested for integration of the MADM cassette by Southern blot. **(D)** Blastocyst injection of targeted ES cells. Targeted ES cells were injected into wild-type blastocysts, allowed to further develop, and embryos were injected into pseudo pregnant dams. F1 chimeric offspring was bred with wild-type animals to obtain germline transmission of the MADM transgenes. **(E)** Generation of experimental MADM mice. In order to generate experimental MADM animals, homozygous stocks for the MADM cassettes in combination with various Cre driver lines were generated first. A typical breeding example generating MADM experimental animals would be to breed a male *MADM*^*TG/TG*^ with a female *MADM*^*GT/GT*^;*Driver*^*Cre/+*^ in order to obtain *MADM*^*GT/TG*^*; Driver*^*Cre/+*^ mice for analysis.

**Figure S4.**
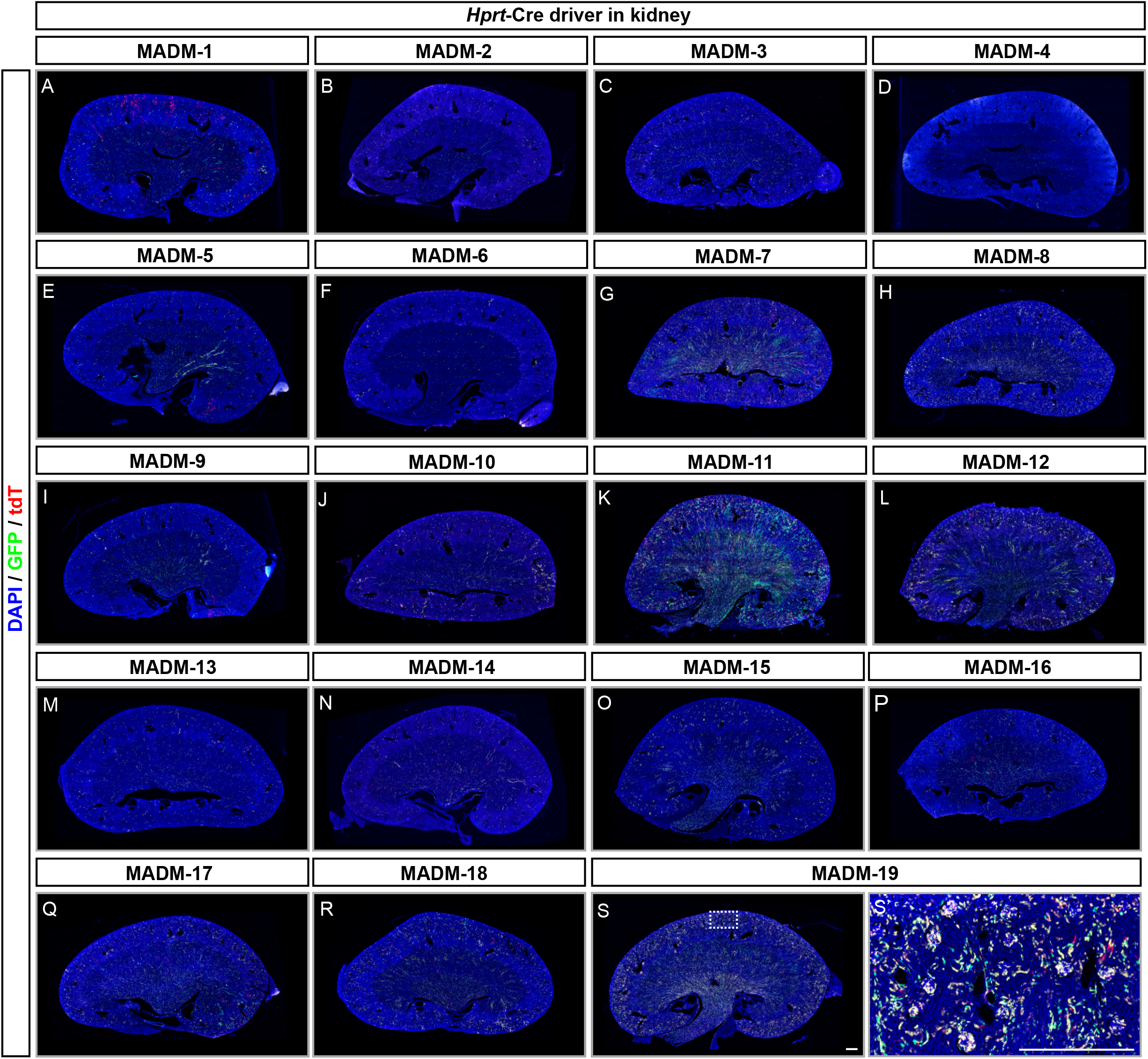
MADM Labeling Pattern in Kidney. **(A-S)** Representative images of sagittal kidney cryosections with MADM labeling (GFP, green; tdT, red) in MADM-1 (A) to MADM-19 (S) in combination with *Hprt*-Cre driver at P21. Inset in S depicts higher resolution image of cortex area (S’). Nuclei are stained by using DAPI (blue). Scale bar: 500μm.

**Figure S5.**
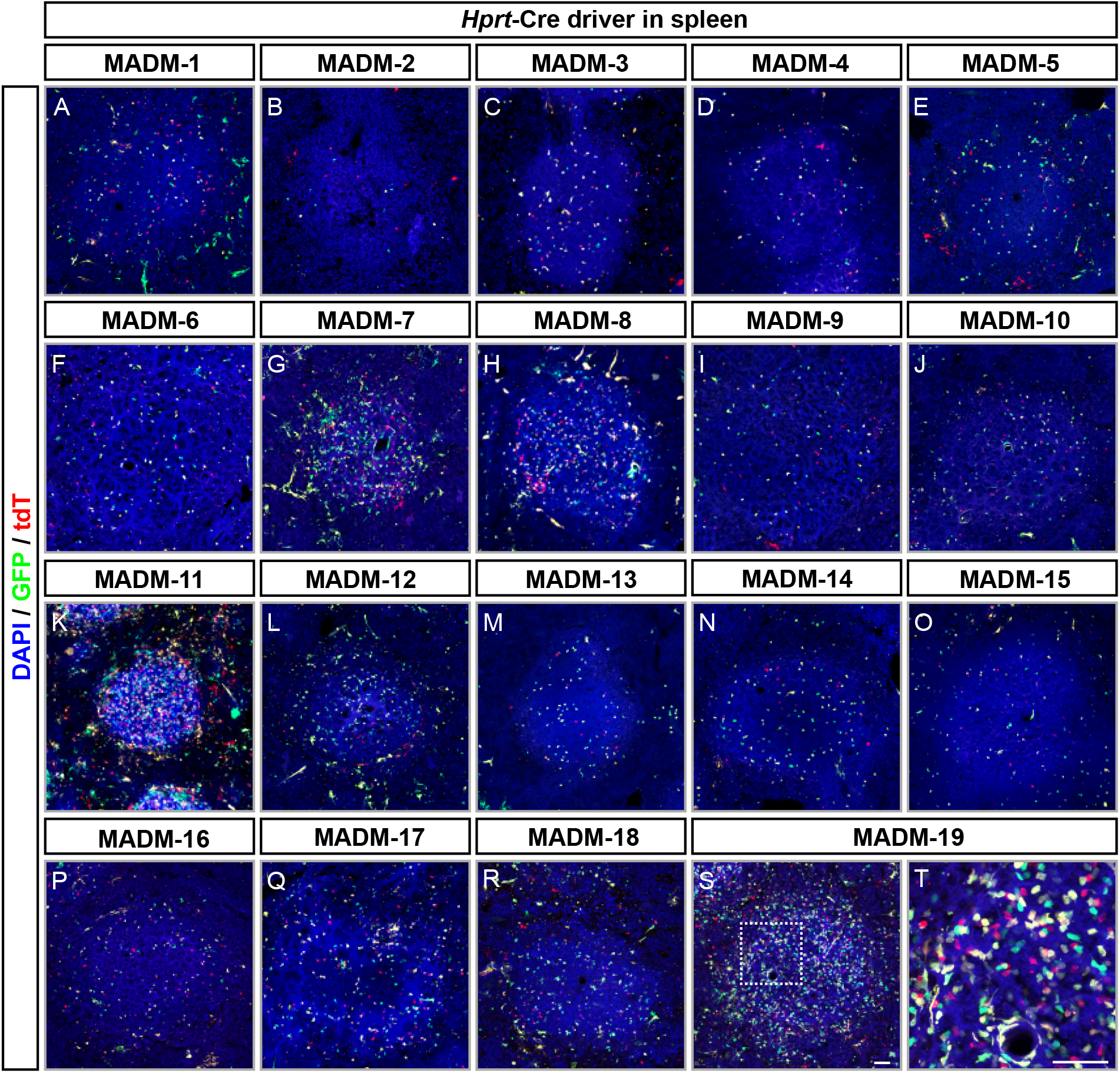
MADM Labeling Pattern in Spleen. **(A-S)** Representative images of horizontal spleen cryosections with MADM labeling (GFP, green; tdT, red) in MADM-1 (A) to MADM-19 (S) in combination with *Hprt*-Cre driver at P21. Areas of white pulp are in the center of the microscopic images. Inset in S shows higher resolution image (S’) of white pulp. Nuclei are stained by using DAPI (blue). Scale bar: 50μm.

**Figure S6.**
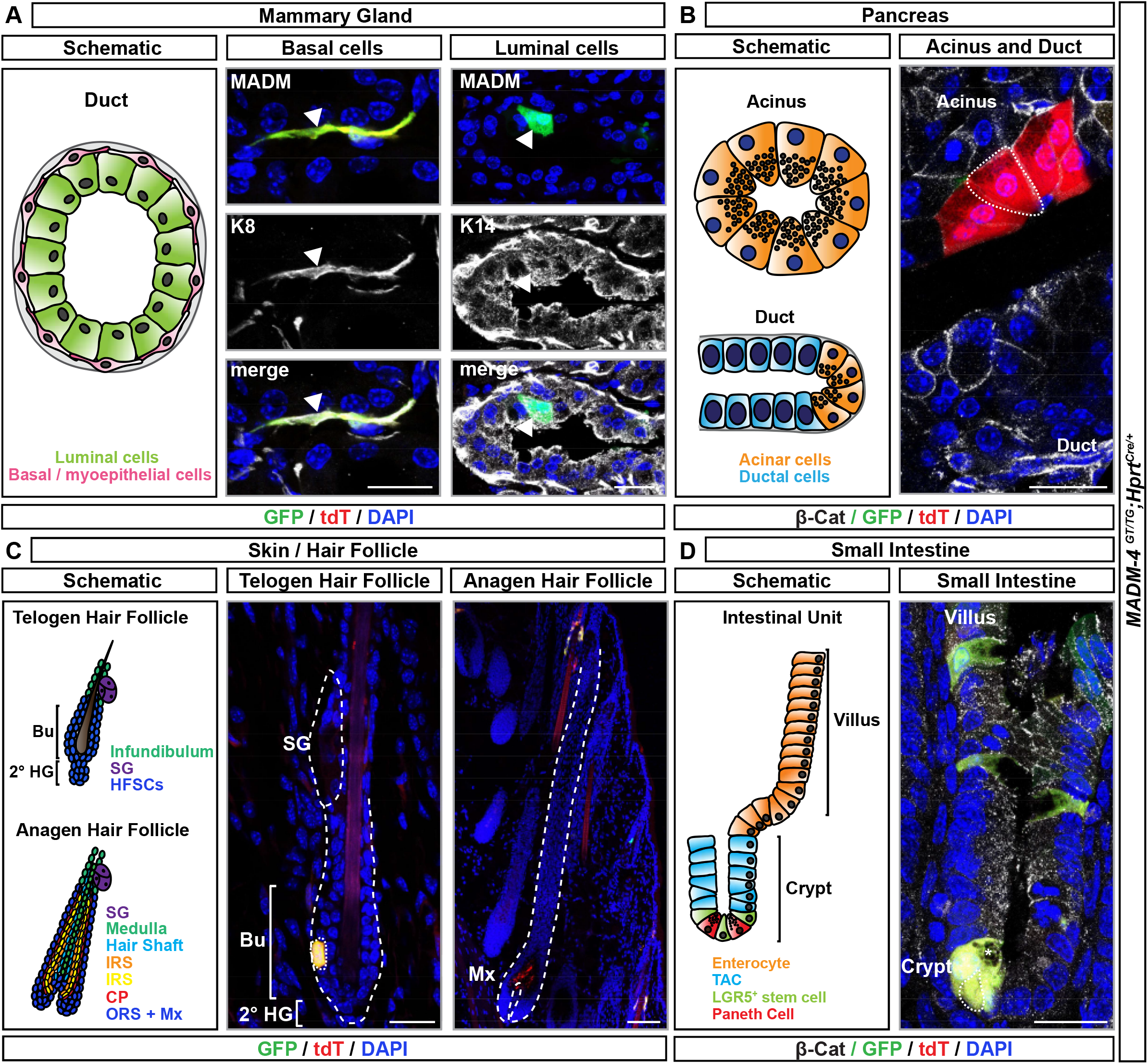
MADM Labeling Pattern in Different Stem Cell Niches. **(A)** Schematic (left) and MADM labeling (middle/right; green, GFP; red, tdT; yellow, GFP/tdT) in mammary gland of lactating *MADM-4*^*GT/TG*^*;Hprt-Cre* female at four months of age. Basal/myoepithelial (middle) and luminal (right) cells are stained with antibodies against K14 and K8 (white), respectively. **(B)** Schematic (left) MADM labeling (right; green, GFP; red, tdT; yellow, GFP/tdT) in *MADM-4*^*GT/TG*^*;Hprt-Cre* pancreas, acinus and duct, at P21. Epithelial cells are visualized by antibody staining against β-Catenin (white, β-Cat). Acinar cells are identified by the presence of intracellular secretory granules. **(C)** Schematic (left) and MADM labeling (middle/right; green, GFP; red, tdT; yellow, GFP/tdT) in telogen (middle) and anagen (right) hair follicles in *MADM-4*^*GT/TG*^*;Hprt-Cre* at P21 (telogen) and P28 (anagen). Bu, bulge; 2° HG, secondary hair germ; SG, sebaceous gland; IRS, inner root sheath; CP, companion layer; ORS, outer root sheath; Mx, matrix. **(D)** Schematic (left) and MADM labeling (right; green, GFP; red, tdT; yellow, GFP/tdT) in small intestine in *MADM-4*^*GT/TG*^*;Hprt-Cre* at P21. Epithelial cells are visualized by antibody staining against β-Catenin (white, β-Cat). Asterick marks a Paneth cell, identified by the presence of intracellular granules. TAC, transit-amplifying cell; LGR5, leucine-rich repeat-containing G-protein coupled receptor 5. Nuclei are stained using DAPI. Scale bar: 20μm.

**Figure S7.**
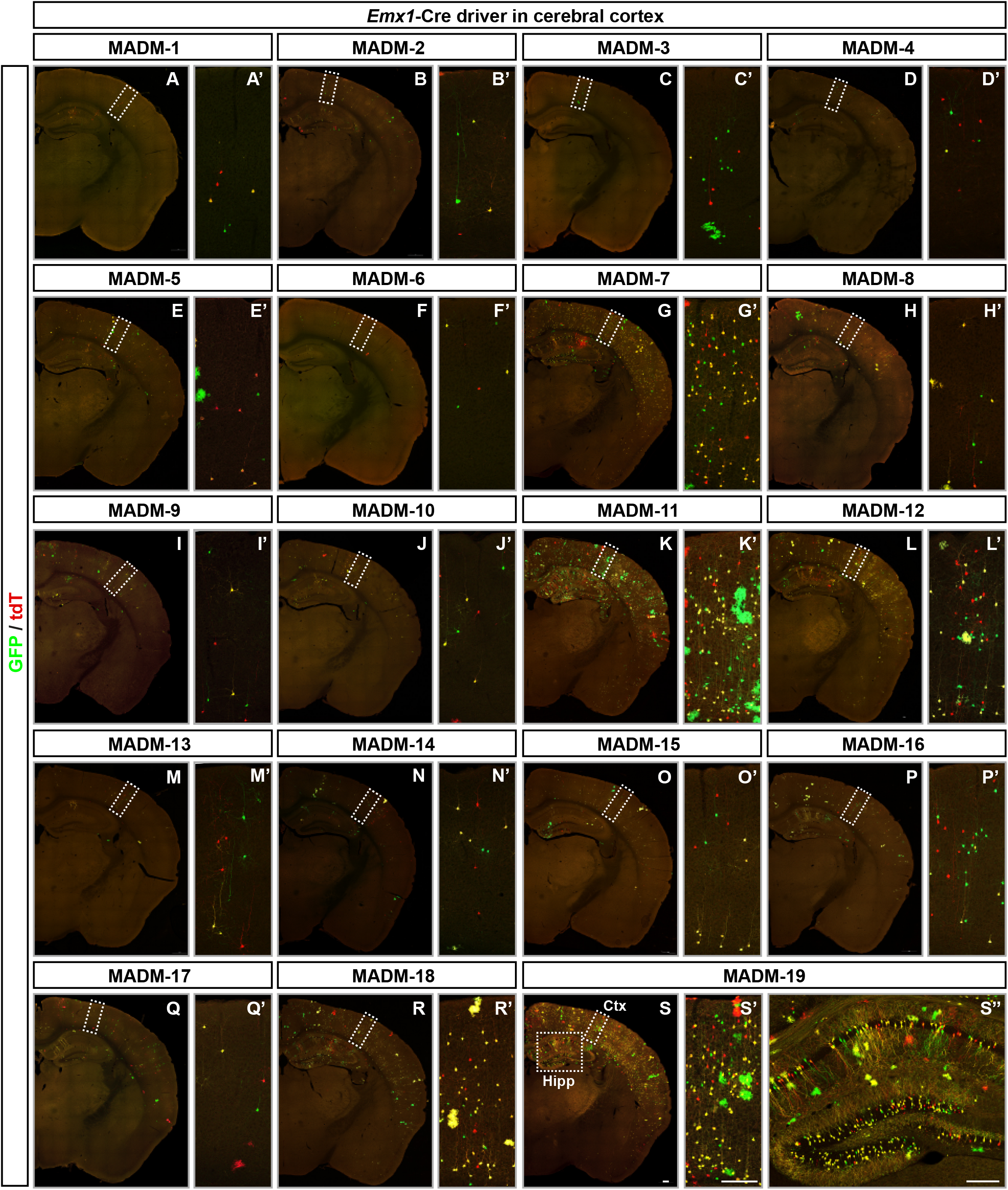
MADM Labeling Pattern in Cerebral Cortex. **(A-S)** Representative images of coronal brain cryosections with MADM labeling (GFP, green; tdT, red) in MADM-1 (A) to MADM-19 (S) in combination with *Emx1*-Cre driver at P21. Insets in S show higher resolution images in neocortex (S’, Ctx) and hippocampus (S’’, Hipp). Scale bar: 200μm.

## Supplemental Tables

**Table S1. Raw Data Obtained from all Experiments.**

Raw data and statistical assessment for charts and panels depicted in Figures 3, 5 and 6.

## STAR METHODS

### KEY RESOURCES TABLE

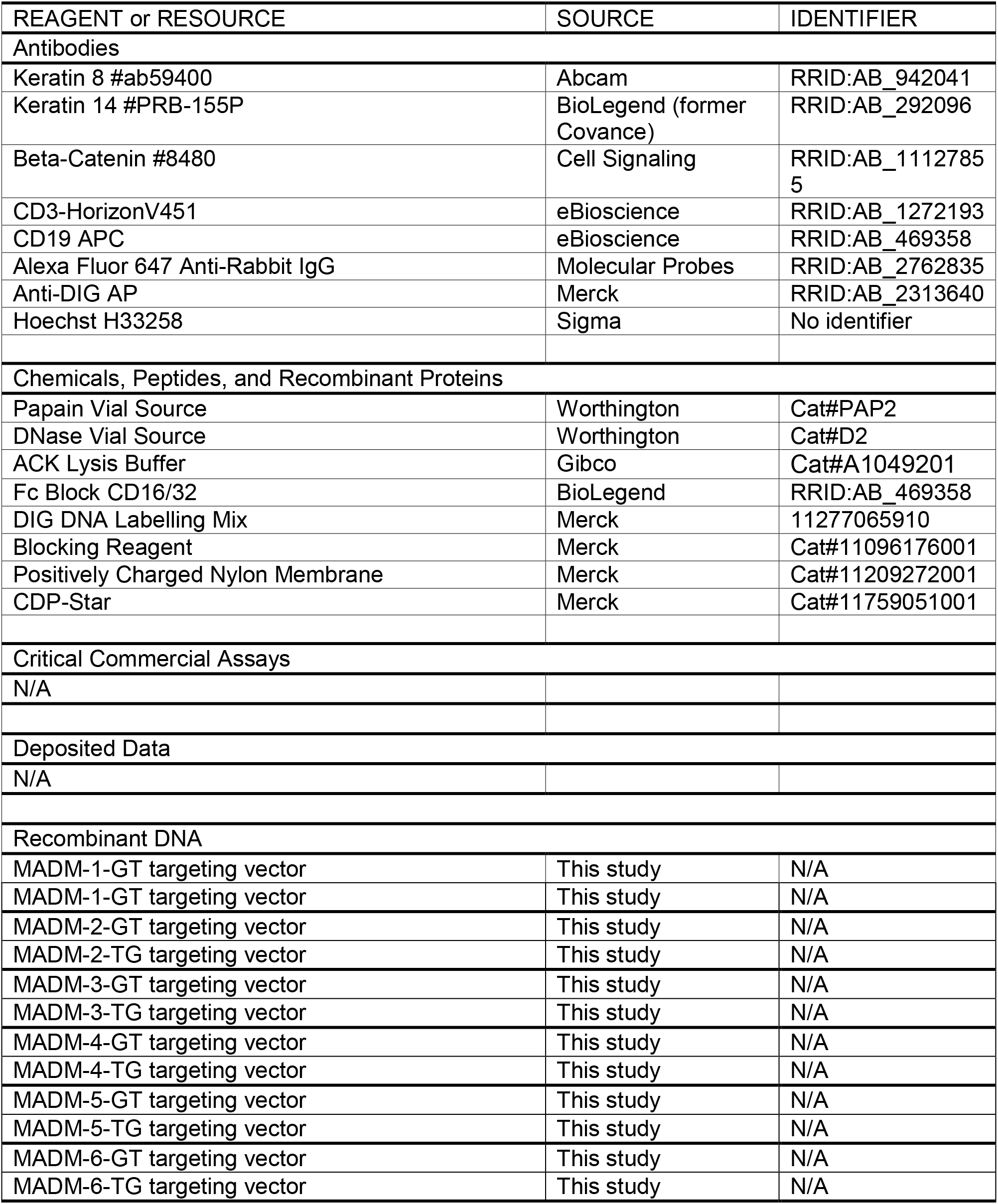
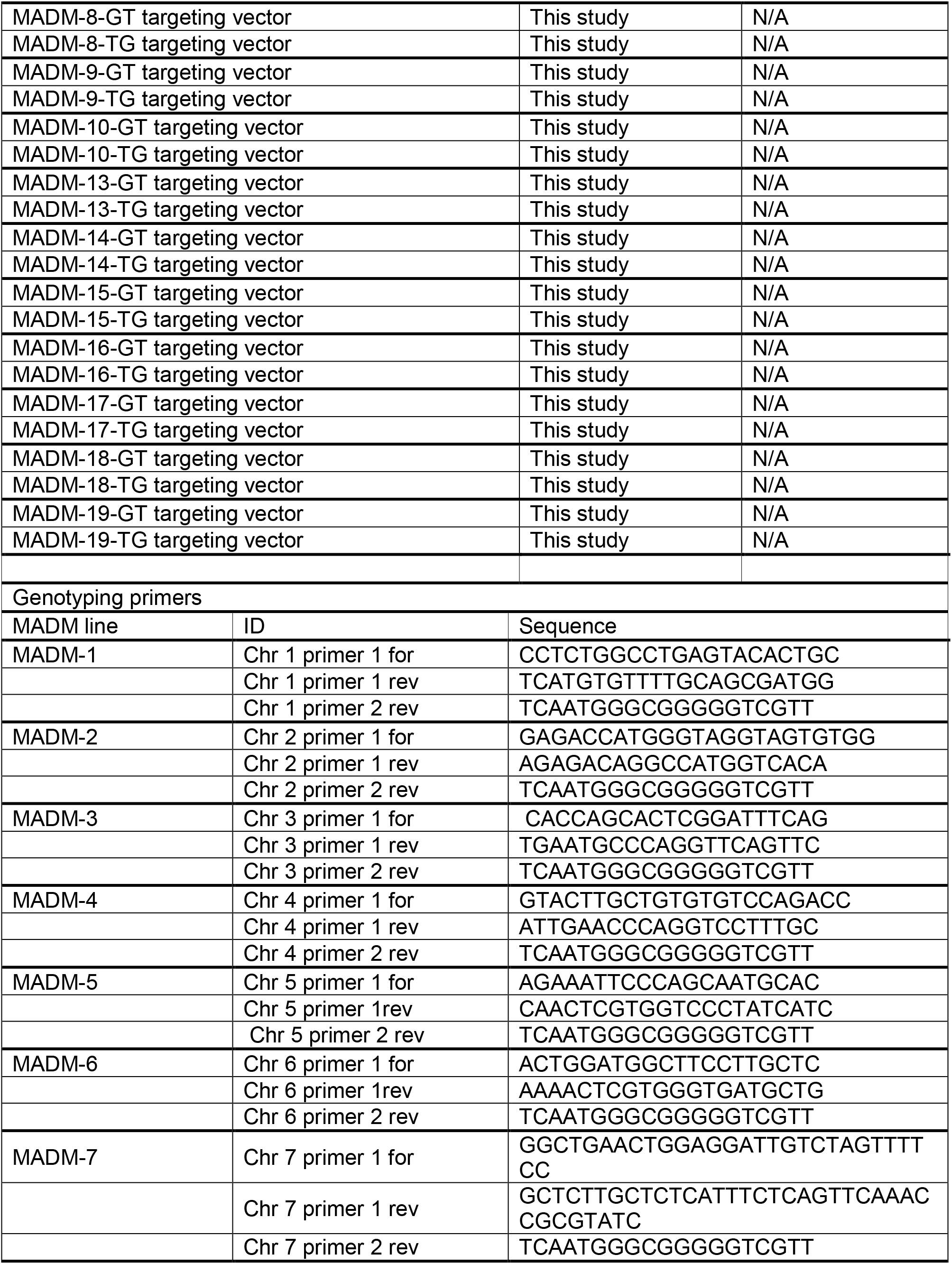
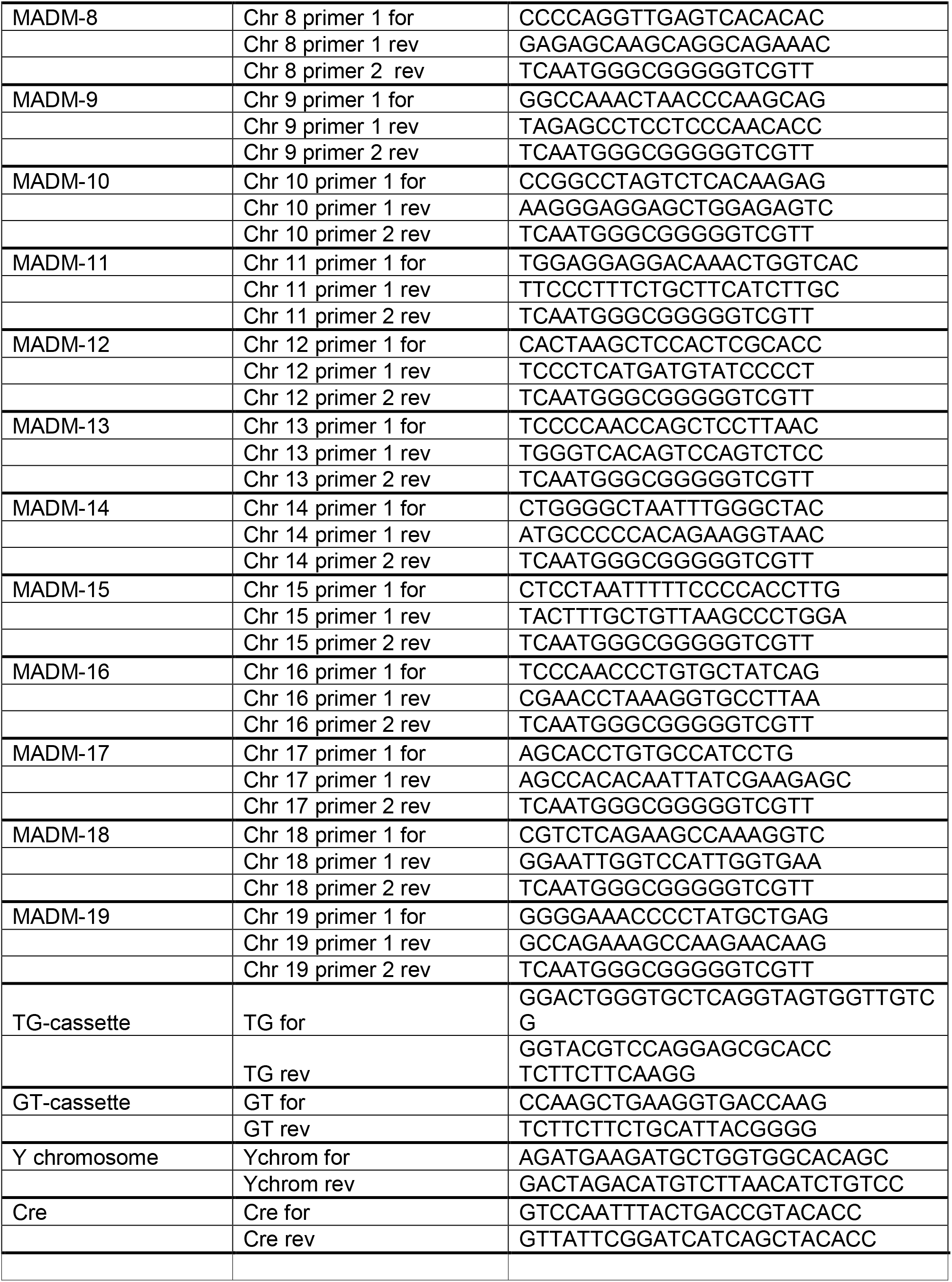
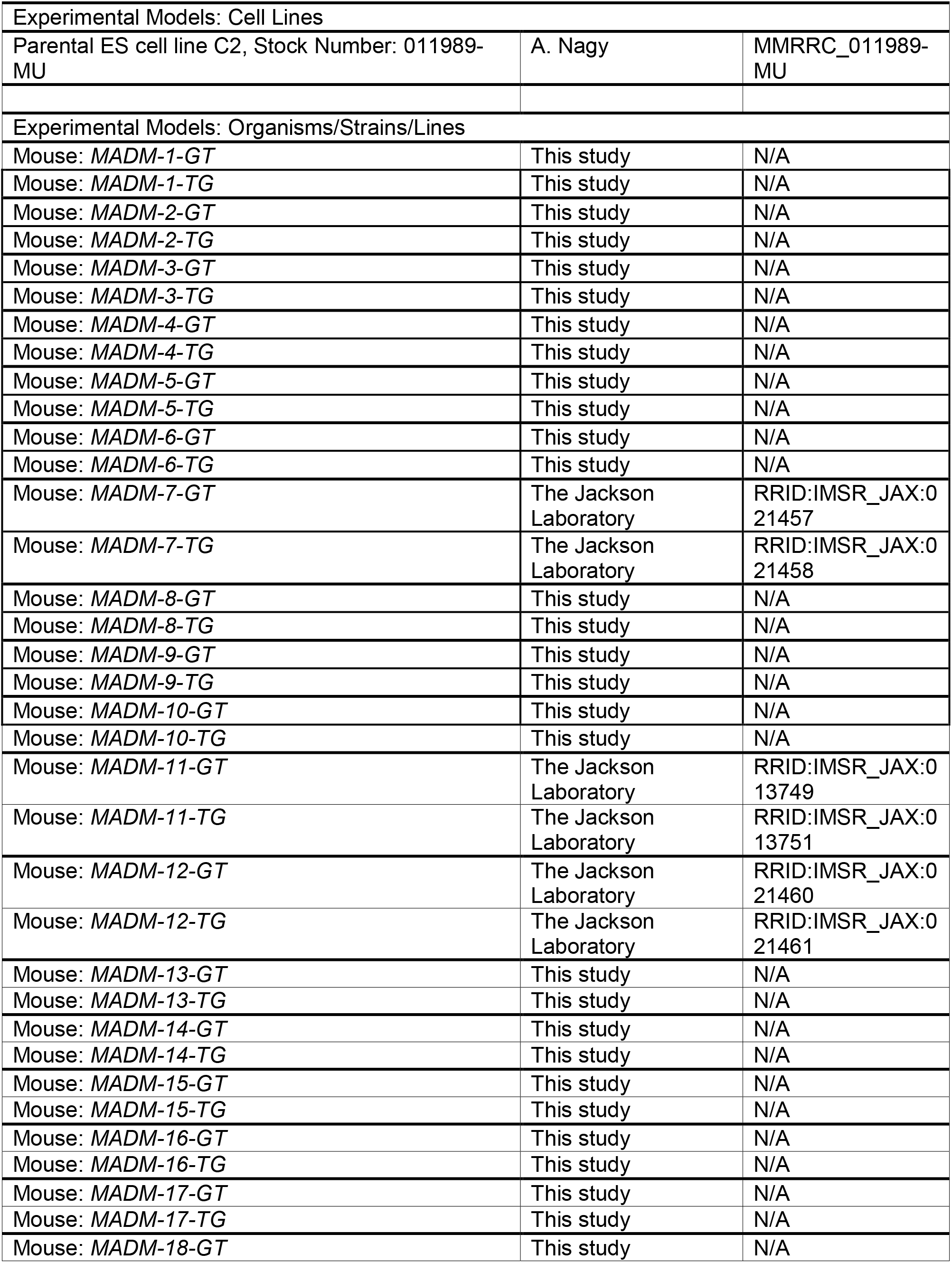
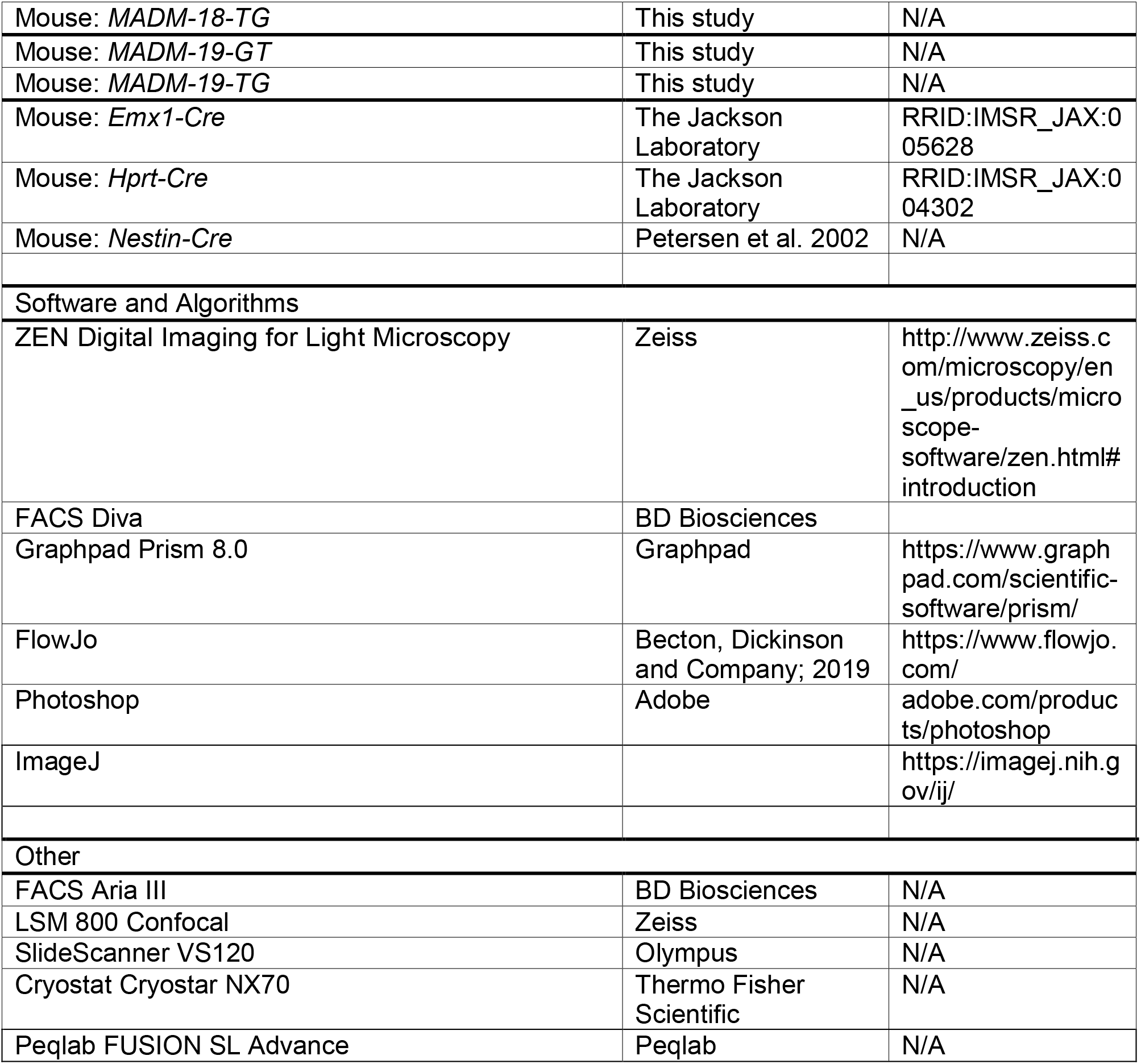

## LEAD CONTACT AND MATERIALS AVAILABILITY

Further information and requests for resources and reagents should be directed to and will be fulfilled by the Lead Contact, Simon Hippenmeyer.

## EXPERIMENTAL MODEL AND SUBJECT DETAILS

### Generation, Breeding and Husbandry of Mouse Lines

Experimental procedures were discussed and approved by the institutional ethics and animal welfare committees at IST Austria, Stanford University, and at University of Veterinary Medicine Vienna in accordance with good scientific practice guidelines and national legislation (license number: IST Austria: BMWF-66.018/0007-II/3b/2012 and BMWFW-66.018/0006-WF/V/3b/2017; University of Veterinary Medicine Vienna: BMWF-68.205/0023-II/3b/2014 and BMBWF-68.205/0010-V/3b/2019). Mice with specific pathogen free status according to FELASA recommendations (Mahler Convenor et al. 2014) were bred and maintained in experimental rodent facilities (room temperature 21 ± 1°C [mean ± SEM]; relative humidity 40%-55%; photoperiod 12L:12D). Food (V1126, Ssniff Spezialitäten GmbH, Soest, Germany) and tap water were available *ad libitum*.

Mouse lines with MADM cassettes inserted in Chr. 7 (Hippenmeyer et al. 2013), Chr. 11 (Hippenmeyer et al. 2010), and Chr. 12 (Hippenmeyer et al. 2013), *Emx1*-Cre (Gorski et al. 2003), *Nestin*-Cre (Petersen et al. 2002), *Hprt*-Cre (Tang et al. 2002) have been described previously. All analyses were carried out in a mixed genetic background. The two lines of each chromosome, with the exception of Chr. 7, 11 and 12, were designated as C57BL/6N;CD1-MADM-GT^tm1(Chr1)Biat^ and C57BL/6N;CD1-MADM-TG^tm1(Chr1)^, as indicated here for Chr. 1. No sex specific differences were observed under any experimental conditions or in any genotype.

## METHOD DETAILS

### Molecular Biology

#### Generation of MADM Targeting Constructs

Molecular cloning and generation of recombinant DNA to construct all plasmids (incl. targeting vectors, plasmids with southern probes etc.), and all nucleic acid procedures as described below were carried out according to standard cloning protocols (Sambrook et al. 1989).

#### Genomic DNA Isolation from Mouse ES Cells

Mouse ES cells were lysed in Lysis Buffer (1M Tris-HCl pH=7.5, 0.5M EDTA, 5M NaCl, 20% Sarcosyl, 20 mg/ml Proteinase K) overnight at 55°C. Next day, the DNA was precipitated with isopropanol for 2 hrs at room temperature with agitation and then carefully transferred into a fresh tube containing TE-buffer. The lids of the tubes were left open for 10 min to allow residual isopropanol to evaporate. The DNA was then incubated for 3 hrs at 37°C.

#### Southern Blot

DIG-labelled probes were generated via PCR amplification of plasmid templates containing the probe sequence using a mix of nucleotides containing Digoxigenin-11-dUTP (DIG-dUTP). The PCR reaction was next separated by electrophoresis and the corresponding band was cut and gel purified using the Monarch DNA gel extraction Kit-NEB.

Genomic DNA was digested with the corresponding enzymes overnight at 37°C and electrophoresed in 0.8% agarose gels for 6 hrs at low voltage together with Lambda Hind III marker. Next day, the agarose gels were depurinized in 0.25M HCl, denaturated in 0.4 NaOH and transferred overnight into a positively charged nylon membrane. Next day, the agarose gels were assessed under UV light to verify complete transfer of DNA to the membrane. The nylon membrane was then neutralized in 0.5M Tris-HCl (pH=7.5) and crosslinked with UV light. The membrane was incubated in hybridization buffer (5x SSC, 2% Blocking reagent, 50% Formamide, 0.1% Sarcosyl, 0.02% SDS) for 4 hrs at 42°C in glass tubes in a rotating oven. In the meantime, the DIG-labelled probe was denaturated at 95°C for 10 min and then quickly chilled on ice for 5 min. The DIG-labeled probe in Hybridization buffer was added to the membrane and incubated overnight at 42°C in glass tubes in a rotating oven. Next day, stringency washes were performed with Wash Solution I (2xSSC, 0.1% SDS) at room temperature, followed by Wash Solution II (0.2x SSC, 0.1% SDS) at 68°C. Next day, the membrane was blocked in blocking solution (1% blocking reagent, 0.1M Maleic acid, 0.15M NaCl) for 1 hr. Then anti-DIG AP antibody (1:20,000) in Blocking Solution was added to the membrane, incubated for 30 min at room temperature and then washed with Wash buffer (0.1M Maleic acid, 0.15M NaCl, 0.3% Tween) for 15 min. Finally the membrane was incubated with CDP-Star (1:100) chemiluminescent substrate in CDP-Star detection buffer (0.1M Tris-HCl, 0.1M NaCl, pH=9.5) for 5 min, wrapped in transparent film and kept in the dark for 1 hr. The pattern of probe hybridization was detected in a Peqlab FUSION SL Advance system for chemiluminescent imaging.

### Generation of Transgenic MADM Mice

#### Targeting of MADM Constructs to Mouse ES Cells by Electroporation

The linearized MADM targeting constructs were introduced into C57BL/6N embryonic stem cells (Parental ES cell line C2, Stock Number: 011989-MU, Citation ID: RRID: MMRRC_011989-MU, A. Nagy Basic ES Cell line) by electroporation using a Bio-Rad Gene Pulser Xcell. After selection with 150μg/ml G418, surviving clones were analyzed for correct targeted integration by southern blot hybridization (see above). Metaphase spread chromosome counting was performed on ES cells of clones with confirmed correct targeting of the MADM cassettes before they were prepared for blastocyst injection.

#### Production of Chimeras

Host blastocysts were produced by superovulation of BALB/cRj females by intraperitoneal (IP) injection with 5.0 IU of equine chorionic gonadotropin (Folligon; Intervet) and, 48 hrs later, with 5.0 IU of human chorionic gonadotropin (Chorulon; Intervet) followed by mating with males of the same strain. Morula stages were harvested from isolated oviducts at day 2.5 days post coitum (dpc) and cultured in M16 medium overnight in an incubator at 37°C and 5% CO_2_ to produce host blastocysts. About 10-15 ES cells were injected into a single blastocyst. The injected embryos were cultured for 2-3 hrs to recover and then transferred into the right uterus horn of 2.5 dpc pseudopregnant RjOrl:Swiss surrogate mothers as described earlier in detail (Rulicke 2004; Rulicke et al. 2006). The offspring were selected based on their chimeric coat color. High-percentage male chimeras (>80%) were bred with C57BL/6NRj females and the offspring were selected by coat color and genotyped by PCR for the respective GT or TG MADM transgenes.

### Genotyping of MADM Reporters

For primer sequences see Key Resources Table. Forward and reverse primer 1 is specific for each MADM reporter. In the absence of MADM cassettes the forward/reverse primer 1 PCR will result in the WT band as indicated. The reverse primer 2 is generic and located in the MADM cassette. The forward/reverse primer 2 PCR will result in the MADM band as indicated. The combined use of all three (forward, reverse primer 1, and reverse primer 2) in a single PCR reaction will enable the distinction of WT (single band at WT size), heterozygote (two bands, one at WT and one at MADM size), and homozygous MADM (single band at MADM size) stock mice. Note that *MADM*^*GT/GT*^ and *MADM*^*TG/TG*^ stock mice should be maintained individually. The distinction of MADM-GT versus MADM-TG is possible by using GT-cassette (GT-for and GT-rev) and TG-cassette (TG for and TG rev) specific primers, respectively. Male mice can be identified by using Y chromosome (Ychrom for and Ychrom rev) specific primers. Presence of transgenes encoding Cre recombinase can be confirmed by using Cre primers (Cre for and Cre rev) as indicated.

### Isolation of MADM-Labeled Tissue

Mice were deeply anesthetized through injection of a ketamine/xylazine/acepromazine solution (65 mg, 13 mg and 2 mg/kg body weight, respectively), and confirmed to be unresponsive through pinching the paw. Perfusion was performed with PBS followed by ice-cold 4% PFA. Tissue was further fixed in 4% PFA overnight at 4°C. Brain, thymus, heart, lung, liver, kidney, spleen, eye and spinal cord were surgically removed and cryopreserved in 30% sucrose for approximately 48 hrs and then embedded in Tissue-Tek O.C.T. (Sakura). All samples were stored at −20°C or −80°C until further usage. Samples were sectioned in a cryo microtome at a 45μm thickness. Brain samples were collected in 24 multi-well dishes and then mounted onto Superfrost Glass Slides (Thermo Fisher Scientific), all other samples were directly mounted on glass slides.

For isolation of skin, pancreas, mammary gland and intestine, no perfusion was required. Mice were sacrificed by cervical dislocation and back skin was prepared for histology as previously described (Amberg et al. 2015). Briefly, back skin was shaved and surgically removed above the spine and placed on lint-free surface. Abdominal mammary glands, pancreas and small intestines were surgically removed. Small intestines were cut open longitudinally and made into Swiss rolls. All samples were incubated in 4% PFA at room temperature for 4hrs, then cryoprotected in 30% sucrose overnight at 4°C and embedded into Tissue-Tek O.C.T. (Sakura). All samples were stored at −20°C. Samples were sectioned at a 20μm thickness and directly mounted onto Superfrost Glass Slides (Thermo Fisher Scientific).

### Histology and Immunostaining of MADM-Labeled Tissue

For immunofluorescence staining in skin, pancreas, mammary gland and intestine, sections were thawed at room temperature for 15 min and encircled with DAKO hydrophobic pen. Then, they were washed 3x for 5 min with PBS. Antigen retrieval was performed by adding pre-warmed citrate buffer pH=6.0 to the samples and incubating them at 85°C for 30 min. Samples were washed 3x for 5 min with PBS. Samples were incubated in blocking solution (10% horse serum, 0.5% Triton X-100 in PBS) for 1h at room temperature. Primary antibodies were diluted in staining solution (5% horse serum, 0.5% Triton X-100 in PBS) and added to the samples over night at 4°C. Next day, the samples were washed 3x for 5min with PBS and incubated with secondary antibodies (1:1000) and Hoechst (Sigma, 1mg/ml stock, 1:1000) diluted in staining solution for 2hrs at room temperature. After washing 3x for 5min with PBS, samples were mounted with Mowiol and stored at 4°C until they were imaged at a Zeiss LSM800. Primary antibodies: Keratin 8 (Abcam), Keratin 14 (BioLegend), beta-Catenin (Cell Signaling). Secondary antibody: donkey anti-rabbit Alexa647 (Molecular Probes). Mounted sections were washed 3x for 5 min in PBS, DAPI stained (1:20’000) for 10 min and then embedded in mounting medium containing 1,4-diazabicyclooctane (DABCO; Roth) and Mowiol (Roth).

### Flow Cytometry

Mice were sacrificed by cervical dislocation and spleens were collected in ice-cold PBS. Spleens were then placed on a 70μm cell strainer on top of a 50ml Falcon tube and minced through the strainer. The strainers were flushed with 10ml PBS-FBS (1x PBS, 2% FBS) and cell suspensions were centrifuged for 6min at 1,200rpm. Cell pellets were resuspended in 1ml ACK lysis buffer (Gibco) and incubated for 30sec. Lysis reaction was stopped by adding 10ml PBS-FBS. The cell suspension was transferred to a 15ml Falcon tube for better visibility of pellet. Cells were centrifuged for 6 min at 1,200 rpm. Pellets were resuspended in 1ml PBS-FBS and transferred to 5ml round-bottom FACS tube via a 70μl cell strainer. Tubes were filled up with PBS-FBS and centrifuged for 6 min at 1,200 rpm. Cells were incubated with Fc block (BD Biosciences) for 5 min and then incubated with 100μl of antibody mastermix for 30min on ice. Antibodies CD3 HorizonV451 (eBioscience) and CD19 APC (eBioscience) were diluted 1:200. Finally, 4ml of PBS-FBS were added and cells were centrifuged for 6 min at 1,200 rpm. Flow cytometric sorting of GFP^+^, tdT^+^ and GFP^+^ tdT^+^ cells was performed on a BD AriaIII. Analysis was performed using FlowJo.

### Analysis of MADM-Labeled Brains and Peripheral Tissue

Representative images were acquired using either an inverted LSM800 or LSM880 with airy scan confocal microscope (Zeiss) and processed using Zeiss Zen Blue software and Photoshop (Adobe). Images for quantification were acquired using a SlideScanner VS120 (Olympus) and processed via custumed scripts in ImageJ. Tiled images, encompassing the entire region of interest, were taken for a minimum of 8 brain sections per animal. Images were imported into Photoshop software (Adobe) and the boundaries for the region of interest were traced. MADM-labeled cells were manually counted based on respective marker expression.

## QUANTIFICATION AND STATISTICAL ANALYSIS

See Table S1 for complete information regarding quantifications and statistics used in this study. This table includes all graphed values, including SEMs, p values, and exact values of n. Statistical analysis was performed in the software Prism8 (GraphPad). Evaluation of data was performed by the two-tailed paired Student’s t-test (Figure 5B), paired ratio t-test (Figure 3V), one-way ANOVA (Figure 5E, 6B) or two-way ANOVA (Figure 6A). Data expressed as ratio was log-transformed previously to the statistical test. For Figure 3V and Figure 5B, n was defined as the density of green/red cells per mm^3^ from one animal resulting from the quantification of 4-20 sections. For Figure 5E and 6A-B, n was defined as the YGR index for one animal resulting from the quantification of 20-24 sections (5E and 6A), or from cells sorted from one animal for Figure 6B. The YGRI was defined as the ratio of yellow cells divided by the average of green and red cells.

## RESOURCES, DATA AND SOFTWARE AVAILABILITY

### Resources

All MADM lines will be made publicly available through *The European Mouse Mutant Archive* (EMMA) and distributed from the University of Veterinary Medicine in Vienna or the Institute of Science and Technology Austria in Klosterneuburg.

## REFERENCES

Ali SR, Hippenmeyer S, Saadat LV, Luo L, Weissman IL, Ardehali R (2014) Existing cardiomyocytes generate cardiomyocytes at a low rate after birth in mice. Proceedings of the National Academy of Sciences of the United States of America 111 (24):8850–8855. doi:10.1073/pnas.1408233111

Amberg N, Holcmann M, Glitzner E, Novoszel P, Stulnig G, Sibilia M (2015) Mouse models of nonmelanoma skin cancer. Methods in molecular biology 1267:217–250. doi:10.1007/978-1-4939-2297-0_10

Apte MS, Meller VH (2012) Homologue pairing in flies and mammals: gene regulation when two are involved. Genetics research international 2012:430587. doi:10.1155/2012/430587

Armakolas A, Klar AJ (2006) Cell type regulates selective segregation of mouse chromosome 7 DNA strands in mitosis. Science 311 (5764):1146–1149. doi:10.1126/science.1120519

Armakolas A, Klar AJ (2007) Left-right dynein motor implicated in selective chromatid segregation in mouse cells. Science 315 (5808):100–101. doi:10.1126/science.1129429

Armakolas A, Koutsilieris M, Klar AJ (2010) Discovery of the mitotic selective chromatid segregation phenomenon and its implications for vertebrate development. Current opinion in cell biology 22 (1):81–87. doi:10.1016/j.ceb.2009.11.006

Barker N, Ridgway RA, van Es JH, van de Wetering M, Begthel H, van den Born M, Danenberg E, Clarke AR, Sansom OJ, Clevers H (2009) Crypt stem cells as the cells-of-origin of intestinal cancer. Nature 457 (7229):608–611. doi:10.1038/nature07602

Barker N, van Es JH, Kuipers J, Kujala P, van den Born M, Cozijnsen M, Haegebarth A, Korving J, Begthel H, Peters PJ, Clevers H (2007) Identification of stem cells in small intestine and colon by marker gene Lgr5. Nature 449 (7165):1003–1007. doi:10.1038/nature06196

Barlow DP, Bartolomei MS (2014) Genomic imprinting in mammals. Cold Spring Harbor perspectives in biology 6 (2). doi:10.1101/cshperspect.a018382

Beattie R, Postiglione MP, Burnett LE, Laukoter S, Streicher C, Pauler FM, Xiao G, Klezovitch O, Vasioukhin V, Ghashghaei TH, Hippenmeyer S (2017) Mosaic Analysis with Double Markers Reveals Distinct Sequential Functions of Lgl1 in Neural Stem Cells. Neuron 94 (3):517–533 e513. doi:10.1016/j.neuron.2017.04.012

Bell CD (2005) Is mitotic chromatid segregation random? Histology and histopathology 20 (4):1313–1320. doi:10.14670/HH-20.1313

Beumer KJ, Pimpinelli S, Golic KG (1998) Induced chromosomal exchange directs the segregation of recombinant chromatids in mitosis of Drosophila. Genetics 150 (1):173–188

Biesecker LG, Spinner NB (2013) A genomic view of mosaicism and human disease. Nature reviews Genetics 14 (5):307–320. doi:10.1038/nrg3424

Bonaguidi MA, Wheeler MA, Shapiro JS, Stadel RP, Sun GJ, Ming GL, Song H (2011) In vivo clonal analysis reveals self-renewing and multipotent adult neural stem cell characteristics. Cell 145 (7):1142–1155. doi:10.1016/j.cell.2011.05.024

Bras-Pereira C, Moreno E (2018) Mechanical cell competition. Current opinion in cell biology 51:15–21. doi:10.1016/j.ceb.2017.10.003

Brennand K, Huangfu D, Melton D (2007) All beta cells contribute equally to islet growth and maintenance. PLoS biology 5 (7):e163. doi:10.1371/journal.pbio.0050163

Buchsbaum IY, Cappello S (2019) Neuronal migration in the CNS during development and disease: insights from in vivo and in vitro models. Development 146 (1). doi:10.1242/dev.163766

Buiting K, Williams C, Horsthemke B (2016) Angelman syndrome - insights into a rare neurogenetic disorder. Nature reviews Neurology 12 (10):584–593. doi:10.1038/nrneurol.2016.133

D’Gama AM, Walsh CA (2018) Somatic mosaicism and neurodevelopmental disease. Nature neuroscience 21 (11):1504–1514. doi:10.1038/s41593-018-0257-3

Desgraz R, Herrera PL (2009) Pancreatic neurogenin 3-expressing cells are unipotent islet precursors. Development 136 (21):3567–3574. doi:10.1242/dev.039214

Devine WP, Wythe JD, George M, Koshiba-Takeuchi K, Bruneau BG (2014) Early patterning and specification of cardiac progenitors in gastrulating mesoderm. eLife 3. doi:10.7554/eLife.03848

Ellis SJ, Gomez NC, Levorse J, Mertz AF, Ge Y, Fuchs E (2019) Distinct modes of cell competition shape mammalian tissue morphogenesis. Nature 569 (7757):497–502. doi:10.1038/s41586-019-1199-y

Espinosa JS, Luo L (2008) Timing neurogenesis and differentiation: insights from quantitative clonal analyses of cerebellar granule cells. The Journal of neuroscience : the official journal of the Society for Neuroscience 28 (10):2301–2312. doi:10.1523/JNEUROSCI.5157-07.2008

Espinosa JS, Wheeler DG, Tsien RW, Luo L (2009) Uncoupling dendrite growth and patterning: single-cell knockout analysis of NMDA receptor 2B. Neuron 62 (2):205–217. doi:10.1016/j.neuron.2009.03.006

Feinberg AP (2007) Phenotypic plasticity and the epigenetics of human disease. Nature 447 (7143):433–440. doi:10.1038/nature05919

Ferguson-Smith AC (2011) Genomic imprinting: the emergence of an epigenetic paradigm. Nature reviews Genetics 12 (8):565–575. doi:10.1038/nrg3032

Ferreira RMM, Sancho R, Messal HA, Nye E, Spencer-Dene B, Stone RK, Stamp G, Rosewell I, Quaglia A, Behrens A (2017) Duct- and Acinar-Derived Pancreatic Ductal Adenocarcinomas Show Distinct Tumor Progression and Marker Expression. Cell reports 21 (4):966–978. doi:10.1016/j.celrep.2017.09.093

Fuchs E, Nowak JA (2008) Building epithelial tissues from skin stem cells. Cold Spring Harbor symposia on quantitative biology 73:333–350. doi:10.1101/sqb.2008.73.032

Gao P, Postiglione MP, Krieger TG, Hernandez L, Wang C, Han Z, Streicher C, Papusheva E, Insolera R, Chugh K, Kodish O, Huang K, Simons BD, Luo L, Hippenmeyer S, Shi SH (2014) Deterministic progenitor behavior and unitary production of neurons in the neocortex. Cell 159 (4):775–788. doi:10.1016/j.cell.2014.10.027

Germani F, Bergantinos C, Johnston LA (2018) Mosaic Analysis in Drosophila. Genetics 208 (2):473–490. doi:10.1534/genetics.117.300256

Gonczy P (2008) Mechanisms of asymmetric cell division: flies and worms pave the way. Nature reviews Molecular cell biology 9 (5):355–366. doi:10.1038/nrm2388

Gonzalez PP, Kim J, Galvao RP, Cruickshanks N, Abounader R, Zong H (2018) p53 and NF 1 loss plays distinct but complementary roles in glioma initiation and progression. Glia 66 (5):999–1015. doi:10.1002/glia.23297

Gorski JA, Balogh SA, Wehner JM, Jones KR (2003) Learning deficits in forebrain-restricted brain-derived neurotrophic factor mutant mice. Neuroscience 121 (2):341–354. doi:10.1016/s0306-4522(03)00426-3

Haig D (2004) Genomic imprinting and kinship: how good is the evidence? Annual review of genetics 38:553–585. doi:10.1146/annurev.genet.37.110801.142741

Henderson NT, Le Marchand SJ, Hruska M, Hippenmeyer S, Luo L, Dalva MB (2019) Ephrin-B3 controls excitatory synapse density through cell-cell competition for EphBs. eLife 8. doi:10.7554/eLife.41563

Hippenmeyer S (2013) Dissection of gene function at clonal level using mosaic analysis with double markers. Front Biol 8 (6):557–568. doi:10.1007/s11515-013-1279-6

Hippenmeyer S (2014) Molecular pathways controlling the sequential steps of cortical projection neuron migration. Advances in experimental medicine and biology 800:1–24. doi:10.1007/978-94-007-7687-6_1

Hippenmeyer S, Johnson RL, Luo L (2013) Mosaic analysis with double markers reveals cell-type-specific paternal growth dominance. Cell reports 3 (3):960–967. doi:10.1016/j.celrep.2013.02.002

Hippenmeyer S, Youn YH, Moon HM, Miyamichi K, Zong H, Wynshaw-Boris A, Luo L (2010) Genetic mosaic dissection of Lis1 and Ndel1 in neuronal migration. Neuron 68 (4):695–709. doi:10.1016/j.neuron.2010.09.027

Hotta Y, Benzer S (1970) Genetic dissection of the Drosophila nervous system by means of mosaics. Proceedings of the National Academy of Sciences of the United States of America 67 (3):1156–1163. doi:10.1073/pnas.67.3.1156

Hsu YC, Li L, Fuchs E (2014) Emerging interactions between skin stem cells and their niches. Nature medicine 20 (8):847–856. doi:10.1038/nm.3643

Jayaraman D, Bae BI, Walsh CA (2018) The Genetics of Primary Microcephaly. Annual review of genomics and human genetics 19:177–200. doi:10.1146/annurev-genom-083117-021441

Joo W, Hippenmeyer S, Luo L (2014) Neurodevelopment. Dendrite morphogenesis depends on relative levels of NT-3/TrkC signaling. Science 346 (6209):626–629. doi:10.1126/science.1258996

Kaplan ES, Ramos-Laguna KA, Mihalas AB, Daza RAM, Hevner RF (2017) Neocortical Sox9+ radial glia generate glutamatergic neurons for all layers, but lack discernible evidence of early laminar fate restriction. Neural development 12 (1):14. doi:10.1186/s13064-017-0091-4

Kim GB, Rincon Fernandez Pacheco D, Saxon D, Yang A, Sabet S, Dutra-Clarke M, Levy R, Watkins A, Park H, Abbasi Akhtar A, Linesch PW, Kobritz N, Chandra SS, Grausam K, Ayala-Sarmiento A, Molina J, Sedivakova K, Hoang K, Tsyporin J, Gareau DS, Filbin MG, Bannykh S, Santiskulvong C, Wang Y, Tang J, Suva ML, Chen B, Danielpour M, Breunig JJ (2019) Rapid Generation of Somatic Mouse Mosaics with Locus-Specific, Stably Integrated Transgenic Elements. Cell 179 (1):251–267 e224. doi:10.1016/j.cell.2019.08.013

Knoblich JA (2008) Mechanisms of asymmetric stem cell division. Cell 132 (4):583–597. doi:10.1016/j.cell.2008.02.007

Knouse KA, Lopez KE, Bachofner M, Amon A (2018) Chromosome Segregation Fidelity in Epithelia Requires Tissue Architecture. Cell 175 (1):200–211 e213. doi:10.1016/j.cell.2018.07.042

Koren S, Reavie L, Couto JP, De Silva D, Stadler MB, Roloff T, Britschgi A, Eichlisberger T, Kohler H, Aina O, Cardiff RD, Bentires-Alj M (2015) PIK3CA(H1047R) induces multipotency and multi-lineage mammary tumours. Nature 525 (7567):114–118. doi:10.1038/nature14669

Kumar ME, Bogard PE, Espinoza FH, Menke DB, Kingsley DM, Krasnow MA (2014) Mesenchymal cells. Defining a mesenchymal progenitor niche at single-cell resolution. Science 346 (6211):1258810. doi:10.1126/science.1258810

Laukoter S, Beattie R, Pauler FM, Amberg N, Nakayama KI, Hippenmeyer S (2020) Imprinted Cdkn1c genomic locus cell-autonomously promotes cell survival in cerebral cortex development. Nature communications 11 (1):195. doi:10.1038/s41467-019-14077-2

Lee AYL, Dubois CL, Sarai K, Zarei S, Schaeffer DF, Sander M, Kopp JL (2018) Cell of origin affects tumour development and phenotype in pancreatic ductal adenocarcinoma. Gut. doi:10.1136/gutjnl-2017-314426

Lee T, Luo L (1999) Mosaic analysis with a repressible cell marker for studies of gene function in neuronal morphogenesis. Neuron 22 (3):451–461. doi:10.1016/s0896-6273(00)80701-1

Lee T, Luo L (2001) Mosaic analysis with a repressible cell marker (MARCM) for Drosophila neural development. Trends in neurosciences 24 (5):251–254. doi:10.1016/s0166-2236(00)01791-4

Liang H, Xiao G, Yin H, Hippenmeyer S, Horowitz JM, Ghashghaei HT (2013) Neural development is dependent on the function of specificity protein 2 in cell cycle progression. Development 140 (3):552–561. doi:10.1242/dev.085621

Liu C, Sage JC, Miller MR, Verhaak RG, Hippenmeyer S, Vogel H, Foreman O, Bronson RT, Nishiyama A, Luo L, Zong H (2011) Mosaic analysis with double markers reveals tumor cell of origin in glioma. Cell 146 (2):209–221. doi:10.1016/j.cell.2011.06.014

Liu P, Jenkins NA, Copeland NG (2002) Efficient Cre-loxP-induced mitotic recombination in mouse embryonic stem cells. Nature genetics 30 (1):66–72. doi:10.1038/ng788

Llorca A, Ciceri G, Beattie R, Wong FK, Diana G, Serafeimidou-Pouliou E, Fernandez-Otero M, Streicher C, Arnold SJ, Meyer M, Hippenmeyer S, Maravall M, Marin O (2019) A stochastic framework of neurogenesis underlies the assembly of neocortical cytoarchitecture. eLife 8. doi:10.7554/eLife.51381

Lozano G, Behringer RR (2007) New mouse models of cancer: single-cell knockouts. Proceedings of the National Academy of Sciences of the United States of America 104 (11):4245–4246. doi:10.1073/pnas.0700173104

Luo L (2007) Fly MARCM and mouse MADM: genetic methods of labeling and manipulating single neurons. Brain research reviews 55 (2):220–227. doi:10.1016/j.brainresrev.2007.01.012

Lv X, Ren SQ, Zhang XJ, Shen Z, Ghosh T, Xianyu A, Gao P, Li Z, Lin S, Yu Y, Zhang Q, Groszer M, Shi SH (2019) TBR2 coordinates neurogenesis expansion and precise microcircuit organization via Protocadherin 19 in the mammalian cortex. Nature communications 10 (1):3946. doi:10.1038/s41467-019-11854-x

Madan E, Gogna R, Moreno E (2018) Cell competition in development: information from flies and vertebrates. Current opinion in cell biology 55:150–157. doi:10.1016/j.ceb.2018.08.002

Mahler Convenor M, Berard M, Feinstein R, Gallagher A, Illgen-Wilcke B, Pritchett-Corning K, Raspa M (2014) FELASA recommendations for the health monitoring of mouse, rat, hamster, guinea pig and rabbit colonies in breeding and experimental units. Laboratory animals 48 (3):178–192. doi:10.1177/0023677213516312

Mayer C, Jaglin XH, Cobbs LV, Bandler RC, Streicher C, Cepko CL, Hippenmeyer S, Fishell G (2015) Clonally Related Forebrain Interneurons Disperse Broadly across Both Functional Areas and Structural Boundaries. Neuron 87 (5):989–998. doi:10.1016/j.neuron.2015.07.011

Merino MM, Levayer R, Moreno E (2016) Survival of the Fittest: Essential Roles of Cell Competition in Development, Aging, and Cancer. Trends in cell biology 26 (10):776–788. doi:10.1016/j.tcb.2016.05.009

Mihalas AB, Hevner RF (2018) Clonal analysis reveals laminar fate multipotency and daughter cell apoptosis of mouse cortical intermediate progenitors. Development 145 (17). doi:10.1242/dev.164335

Mohamed TMA, Ang YS, Radzinsky E, Zhou P, Huang Y, Elfenbein A, Foley A, Magnitsky S, Srivastava D (2018) Regulation of Cell Cycle to Stimulate Adult Cardiomyocyte Proliferation and Cardiac Regeneration. Cell 173 (1):104–116 e112. doi:10.1016/j.cell.2018.02.014

Morgan TH, Bridges CB (1919) Contributions to the genetics of Drosophila melanogaster. The origin of gynandromorphs. Publs Carnegie Instn 278:1–122

Muzumdar MD, Dorans KJ, Chung KM, Robbins R, Tammela T, Gocheva V, Li CM, Jacks T (2016) Clonal dynamics following p53 loss of heterozygosity in Kras-driven cancers. Nature communications 7:12685. doi:10.1038/ncomms12685

Muzumdar MD, Luo L, Zong H (2007) Modeling sporadic loss of heterozygosity in mice by using mosaic analysis with double markers (MADM). Proceedings of the National Academy of Sciences of the United States of America 104 (11):4495–4500. doi:10.1073/pnas.0606491104

Ortiz-Alvarez G, Daclin M, Shihavuddin A, Lansade P, Fortoul A, Faucourt M, Clavreul S, Lalioti ME, Taraviras S, Hippenmeyer S, Livet J, Meunier A, Genovesio A, Spassky N (2019) Adult Neural Stem Cells and Multiciliated Ependymal Cells Share a Common Lineage Regulated by the Geminin Family Members. Neuron 102 (1):159–172 e157. doi:10.1016/j.neuron.2019.01.051

Petersen PH, Zou K, Hwang JK, Jan YN, Zhong W (2002) Progenitor cell maintenance requires numb and numblike during mouse neurogenesis. Nature 419 (6910):929–934. doi:10.1038/nature01124

Picco N, Hippenmeyer S, Rodarte J, Streicher C, Molnar Z, Maini PK, Woolley TE (2019) A mathematical insight into cell labelling experiments for clonal analysis. Journal of anatomy 235 (3):687–696. doi:10.1111/joa.13001

Pimpinelli S, Ripoll P (1986) Nonrandom segregation of centromeres following mitotic recombination in Drosophila melanogaster. Proceedings of the National Academy of Sciences of the United States of America 83 (11):3900–3903. doi:10.1073/pnas.83.11.3900

Pinson A, Namba T, Huttner WB (2019) Malformations of Human Neocortex in Development - Their Progenitor Cell Basis and Experimental Model Systems. Frontiers in cellular neuroscience 13:305. doi:10.3389/fncel.2019.00305

Riccio P, Cebrian C, Zong H, Hippenmeyer S, Costantini F (2016) Ret and Etv4 Promote Directed Movements of Progenitor Cells during Renal Branching Morphogenesis. PLoS biology 14 (2):e1002382. doi:10.1371/journal.pbio.1002382

Rossant J, Spence A (1998) Chimeras and mosaics in mouse mutant analysis. Trends in genetics : TIG 14 (9):358–363. doi:10.1016/s0168-9525(98)01552-2

Rulicke T (2004) Pronuclear microinjection of mouse zygotes. Methods in molecular biology 254:165–194. doi:10.1385/1-59259-741-6:165

Rulicke T, Haenggli A, Rappold K, Moehrlen U, Stallmach T (2006) No transuterine migration of fertilised ova after unilateral embryo transfer in mice. Reproduction, fertility, and development 18 (8):885–891. doi:10.1071/rd06054

Salpeter SJ, Klein AM, Huangfu D, Grimsby J, Dor Y (2010) Glucose and aging control the quiescence period that follows pancreatic beta cell replication. Development 137 (19):3205–3213. doi:10.1242/dev.054304

Sambrook J, Fritsch EF, Maniatis T (1989) Molecular Cloning: a laboratory manual. Cold Spring Harbor Laboratory Press.

Shi W, Xianyu A, Han Z, Tang X, Li Z, Zhong H, Mao T, Huang K, Shi SH (2017) Ontogenetic establishment of order-specific nuclear organization in the mammalian thalamus. Nature neuroscience 20 (4):516–528. doi:10.1038/nn.4519

Smith FM, Garfield AS, Ward A (2006) Regulation of growth and metabolism by imprinted genes. Cytogenetic and genome research 113 (1-4):279–291. doi:10.1159/000090843

Snippert HJ, van der Flier LG, Sato T, van Es JH, van den Born M, Kroon-Veenboer C, Barker N, Klein AM, van Rheenen J, Simons BD, Clevers H (2010) Intestinal crypt homeostasis results from neutral competition between symmetrically dividing Lgr5 stem cells. Cell 143 (1):134–144. doi:10.1016/j.cell.2010.09.016

Stern C (1936) Somatic Crossing over and Segregation in Drosophila Melanogaster. Genetics 21 (6):625–730

Subramanian L, Calcagnotto ME, Paredes MF (2019) Cortical Malformations: Lessons in Human Brain Development. Frontiers in cellular neuroscience 13:576. doi:10.3389/fncel.2019.00576

Sun Q, Lee W, Mohri Y, Takeo M, Lim CH, Xu X, Myung P, Atit RP, Taketo MM, Moubarak RS, Schober M, Osman I, Gay DL, Saur D, Nishimura EK, Ito M (2019) A novel mouse model demonstrates that oncogenic melanocyte stem cells engender melanoma resembling human disease. Nature communications 10 (1):5023. doi:10.1038/s41467-019-12733-1

Tang SH, Silva FJ, Tsark WM, Mann JR (2002) A Cre/loxP-deleter transgenic line in mouse strain 129S1/SvImJ. Genesis 32 (3):199–202. doi:10.1002/gene.10030

Tasic B, Miyamichi K, Hippenmeyer S, Dani VS, Zeng H, Joo W, Zong H, Chen-Tsai Y, Luo L (2012) Extensions of MADM (mosaic analysis with double markers) in mice. PloS one 7 (3):e33332. doi:10.1371/journal.pone.0033332

Taverna E, Gotz M, Huttner WB (2014) The cell biology of neurogenesis: toward an understanding of the development and evolution of the neocortex. Annual review of cell and developmental biology 30:465–502. doi:10.1146/annurev-cellbio-101011-155801

Tucci V, Isles AR, Kelsey G, Ferguson-Smith AC, Erice Imprinting G (2019) Genomic Imprinting and Physiological Processes in Mammals. Cell 176 (5):952–965. doi:10.1016/j.cell.2019.01.043

Tuna M, Knuutila S, Mills GB (2009) Uniparental disomy in cancer. Trends in molecular medicine 15 (3):120–128. doi:10.1016/j.molmed.2009.01.005

Van Keymeulen A, Lee MY, Ousset M, Brohee S, Rorive S, Giraddi RR, Wuidart A, Bouvencourt G, Dubois C, Salmon I, Sotiriou C, Phillips WA, Blanpain C (2015) Reactivation of multipotency by oncogenic PIK3CA induces breast tumour heterogeneity. Nature 525 (7567):119–123. doi:10.1038/nature14665

Van Keymeulen A, Rocha AS, Ousset M, Beck B, Bouvencourt G, Rock J, Sharma N, Dekoninck S, Blanpain C (2011) Distinct stem cells contribute to mammary gland development and maintenance. Nature 479 (7372):189–193. doi:10.1038/nature10573

Wong SZH, Scott EP, Mu W, Guo X, Borgenheimer E, Freeman M, Ming GL, Wu QF, Song H, Nakagawa Y (2018) In vivo clonal analysis reveals spatiotemporal regulation of thalamic nucleogenesis. PLoS biology 16 (4):e2005211. doi:10.1371/journal.pbio.2005211

Wuidart A, Sifrim A, Fioramonti M, Matsumura S, Brisebarre A, Brown D, Centonze A, Dannau A, Dubois C, Van Keymeulen A, Voet T, Blanpain C (2018) Early lineage segregation of multipotent embryonic mammary gland progenitors. Nature cell biology 20 (6):666–676. doi:10.1038/s41556-018-0095-2

Xu T, Rubin GM (1993) Analysis of genetic mosaics in developing and adult Drosophila tissues. Development 117 (4):1223–1237

Yadlapalli S, Yamashita YM (2013) Chromosome-specific nonrandom sister chromatid segregation during stem-cell division. Nature 498 (7453):251–254. doi:10.1038/nature12106

Yamashita YM (2013) Nonrandom sister chromatid segregation of sex chromosomes in Drosophila male germline stem cells. Chromosome research : an international journal on the molecular, supramolecular and evolutionary aspects of chromosome biology 21 (3):243–254. doi:10.1007/s10577-013-9353-0

Yamazawa K, Ogata T, Ferguson-Smith AC (2010) Uniparental disomy and human disease: an overview. American journal of medical genetics Part C, Seminars in medical genetics 154C (3):329–334. doi:10.1002/ajmg.c.30270

Yao M, Ventura PB, Jiang Y, Rodriguez FJ, Wang L, Perry JSA, Yang Y, Wahl K, Crittenden RB, Bennett ML, Qi L, Gong CC, Li XN, Barres BA, Bender TP, Ravichandran KS, Janes KA, Eberhart CG, Zong H (2020) Astrocytic trans-Differentiation Completes a Multicellular Paracrine Feedback Loop Required for Medulloblastoma Tumor Growth. Cell 180 (3):502–520 e519. doi:10.1016/j.cell.2019.12.024

Yizhak K, Aguet F, Kim J, Hess JM, Kubler K, Grimsby J, Frazer R, Zhang H, Haradhvala NJ, Rosebrock D, Livitz D, Li X, Arich-Landkof E, Shoresh N, Stewart C, Segre AV, Branton PA, Polak P, Ardlie KG, Getz G (2019) RNA sequence analysis reveals macroscopic somatic clonal expansion across normal tissues. Science 364 (6444). doi:10.1126/science.aaw0726

Yochem J, Herman RK (2003) Investigating C. elegans development through mosaic analysis. Development 130 (20):4761–4768. doi:10.1242/dev.00701

Zong H, Espinosa JS, Su HH, Muzumdar MD, Luo L (2005) Mosaic analysis with double markers in mice. Cell 121 (3):479–492. doi:10.1016/j.cell.2005.02.012

Zugates CT, Lee T (2004) Genetic mosaic analysis in the nervous system. Current opinion in neurobiology 14 (5):647–653. doi:10.1016/j.conb.2004.08.005

